# Breaking the redundancy: TAZ outperforms YAP1 in GIST progression

**DOI:** 10.1101/2025.09.02.673653

**Authors:** Irène Pezzati, César Serrano, Eric Vivès, Frédéric Chibon, Sandrine Faure, Pascal de Santa Barbara, Sébastien Deshayes, Prisca Boisguérin

## Abstract

**Background:** Gastrointestinal stromal tumors (GIST) are mainly caused by gain-of-function mutations in *KIT* or *PDGFRA* genes and are the most common neoplasms of the digestive tract. Imatinib (IM), a tyrosine kinase inhibitor (TKI) targeting these oncogenic drivers, has considerably improved patient outcomes, although resistance remains a major challenge. The transcriptional co-activators YAP1 and TAZ, downstream effectors of the Hippo pathway, have emerged as potential oncogenic drivers in various cancers, including GIST. However, their specific roles in KIT-dependent tumor development remain unclear.

**Methods:** We used WRAP5-based nanoparticles loaded with specifically designed siRNA to selectively silence YAP1 and/or TAZ proteins in KIT-dependent IM-sensitive GIST cell lines. This nucleic acid delivery system enabled efficient and specific knockdown without cytotoxicity. We assessed the impact on cell proliferation, migration, and gene expression, focusing on YAP1/TAZ targets *CYR61* and *CTGF*.

**Results:** TAZ silencing resulted in a substantial reduction in GIST-T1 cell proliferation and migration, whereas YAP1 knockdown was comparatively limited. This finding was consistent with an increased *TAZ* expression in GIST patients, which was associated with shorter progression-free survival and an increased tendency for metastasis development. A slight additive effect was observed upon a combined YAP1/TAZ silencing in the migration assay, suggesting a more complex regulation between these two proteins. CYR61 and CTGF expressions were predominantly regulated by TAZ, though a stronger downregulation was observed upon dual knockdown in a subset of GIST cell lines with differential YAP1 and TAZ basal expression. Finally, CYR61 seemed to be more implicated in cell proliferation inhibition, which is further supported by the correlation between high *CYR61* expression and poor prognosis in the GIST patients.

**Conclusion:** Our results highlight the central regulatory function of TAZ-CYR61 axis in oncogenic processes in KIT-dependent GIST, with a modest contribution from YAP1. Targeting TAZ, alone or in combination with YAP1, may represent a promising therapeutic approach, particularly in the context of tumor heterogeneity.

## BACKGROUND

Gastrointestinal stromal tumors (GIST) are the most prevalent mesenchymal cancers of the digestive tract, with an estimated annual incidence of 10-15 per million individuals wordwilde [1,2]. The histogenesis of these rare sarcomas is attributed to interstitial cells of Cajal (ICC), which act as pacemakers regulating gastrointestinal motility, or related mesenchymal progenitors [3]. GIST are primarily driven by oncogenic gain-of-function mutations in receptor tyrosine kinase genes, most notably *KIT* (75-80% of cases) and, to a lesser extent, *PDGFRA* (5-10%) [4,5]. These mutations lead to the constitutive activation of downstream signaling pathways, resulting in uncontrolled cellular proliferation and survival [6,7]. The process of oncogenic signaling cascades downstream of KIT involves the PI3K/AKT/mTOR and RAS/MAPK/ERK pathways. These pathways play critical roles in GIST development and progression, contributing to tumor initiation, growth, and metastatic dissemination [1,8,9].

The introduction of tyrosine kinase inhibitors (TKI) in the early 21^st^ century, such as imatinib mesylate (IM), has significantly improved patient outcomes by effectively inhibiting KIT-downstream PI3K and MAPK pathways [10]. Nevertheless, resistance remains a major clinical challenge in advanced and metastatic GIST, as most patients experience disease progression within 18 to 24 months of treatment, due to secondary resistance mechanisms [7]. Second- and third-line TKI therapies, including sunitinib and regorafenib, have been demonstrated to induce temporary disease control [11,12]. However, their clinical efficacy is constrained by their capacity to target only a subset of secondary *KIT* mutations and by the molecular heterogeneity of resistance [5,6,13]. Another limitation shared by these therapies is their common mechanism of action: they do not address the emergence of additional mutations or the broader cellular adaptations that occur during prolonged TKI exposure [1]. In certain instances, resistance to TKI treatment can result from mechanisms that entirely bypass KIT, such as the loss of KIT expression. This results in a KIT-independent GIST phenotype, which is unresponsive to TKI treatments [14–16]. Together, these limitations underscore the urgent need for alternative therapeutic strategies and a deeper understanding of the molecular pathways that drive GIST progression and resistance.

Recent studies have identified LImb eXpression 1 (LIX1) as a novel contributor of GIST progression and therapeutic resistance. LIX1 is overexpressed in aggressive and relapsed tumors, and its levels increase upon TKI resistance [17,18]. Functionally, LIX1 promotes MAPK pathway reactivation following KIT inhibition, thereby limiting TKI efficacy. LIX1 also regulates mitochondrial function, proliferation, and tumoral lineage identity in GIST cells in KIT-dependent contexts through the Hippo pathway and its two paralogous downstream effectors, the Yes-associated protein (YAP1) and the WW domain-containing transcription regulator protein 1 (WWTR1), also known as TAZ [17–19].

The Hippo signaling pathway is a highly conserved regulator of organ size, cell proliferation, and apoptosis, playing a crucial role in tissue homeostasis [20]. In its active form, the Hippo core function is to restrict YAP1 and TAZ to the cytoplasm by LAST1/2-depending phosphorylation of both proteins [21]. Upon Hippo inactivation, YAP1 and TAZ are dephosphorylated and translocate to the nucleus, where they mainly bind TEAD transcription factors [21]. Together, YAP1/TAZ/TEAD complex drive the expression of genes involved in cell growth, migration, survival, and tissue identity in numerous organs, including the digestive tract, such as *CTGF*, *CYR61*, *AREG, MYC, MCL-1, BIRC5, AXL, etc.* [22–25]. Furthermore, in the nucleus, YAP1 and TAZ also participate in the expression of Cyclin D1 (CD1), a key cell cycle regulator of the G1/S transition implicated in tumor invasion and resistance to therapy in various cancers (pancreatic cancer, non-small cell lung carcinoma, breast cancer, etc.) [26,27].

Focusing on GIST, recent proof-of-concept studies have shown that dysregulation of the Hippo pathway, with YAP1 and TAZ as positive regulators of CD1 can drive tumor growth and progression. These findings establish Hippo/YAP1/CD1 signaling as a key oncogenic mechanism in KIT-independent IM-resistant GIST [14,15]. In KIT-dependent GIST cells, sensitive and resistant to IM, pharmacological inhibition by the Bcl-2 inhibitor, AT101, of the YAP1/TAZ-CD1 axe has been recently shown to reduce tumor growth and induce apoptosis in preclinical GIST models [28]. Moreover, pharmacological inhibition of YAP1/TAZ with verteporfin promotes ferroptosis in KIT-dependent IM-sensitive and -resistant human GIST cells. This result correlates with aggressive tumor behavior, underscoring its therapeutic potential in cases resistant to TKI [29,30].

It has been reported that the Hippo pathway is often dysregulated in combination with hyperactivation of YAP1/TAZ in various human cancers, including solid tumors and sarcomas [31]. In these cases, dysregulation contributes to aggressive phenotypes, therapy resistance, and metastatic dissemination [32,33]. Despite their frequent co-activation in cancers, the term “YAP1/TAZ” is often described as interchangeable transcriptional co-activators within the Hippo signaling pathway and studied as a single unit without any distinction in functional assays [34,35]. Yet accumulating evidence reveals fundamental differences between them, not only in their protein domain composition and transcriptional partners, but also in their patterns of expression and regulatory behavior, which vary depending on tissue context and physiological conditions [36,37]. These distinctions call for a careful evaluation of their respective roles in cancer, especially in sarcomas [38–41].

Based on these facts, we hypothesized that YAP1 and TAZ could be promising therapeutic targets for GIST. Therefore, we aimed to individually and jointly evaluate the contribution of YAP1 and TAZ in KIT-dependent IM-sensitive GIST cells using specifically designed small interfering RNA (siRNA). For cellular internalization, we opted for the tryptophan and arginine-rich amphipathic peptide, WRAP5, which self-assembled into peptide-based nanoparticles (PBN) in the presence of siRNA as payload [42]. Formulated at a specific molar ratio (WRAP5:siRNA = 20:1), WRAP5-based nanoparticles were selected from a performed structure-activity study as one of the lead peptides [43], which was mainly evaluated in the U87 glioblastoma cell line [42]. More recently, we have demonstrated that WRAP5 could encapsulate up to three distinct siRNA, enabling the simultaneous silencing of three different proteins within GIST cells [44]. Compared to conventional lipid-based transfection reagents, the PBN demonstrated superior safety profiles with minimal cytotoxicity in long-term assays, while maintaining high intracellular delivery efficiency [43,45]. Its versatility has been validated across multiple cell types, both cancerous and non-cancerous, as well as in animal models (mouse and zebrafish), making it particularly suitable for studying dose-dependent gene silencing effects in therapeutic contexts as exemplified through different protein silencing [44,46–48].

In the present study, the efficacy of YAP1 and/or TAZ silencing was firstly confirmed in human KIT-dependent IM-sensitive GIST-T1 cells. For the first time, we could demonstrate that TAZ is a regulator of proliferation and migration in GIST-T1 cells, with a limited contribution of YAP1. This predominant role was in accordance with a bad prognosis in GIST patients with high level of *TAZ* that we revealed from the ATGsarc database. Consequently, we demonstrated that this phenomenon is attributable to the transcriptional activity of TAZ on *CTGF* and *CYR61* genes. Finally, the role of TAZ, and a slight additive effect of combined YAP1 and TAZ was also validated in other GIST cell lines (GIST-430 and GIST-882). In summary, the present findings newly underscore the significance of TAZ as a key player in GIST development.

## METHODS

### Peptide

WRAP5 peptide (LLRLLRWWWRLLRLL) was synthesized at the SynBio3 platform (IBMM Montpellier), and the crude product was purified in-house following a qualitative analysis by HPLC/MS (∼95% purity). WRAP5 stock solutions were prepared at 400 µM or 40 µM and stored at 4°C.

### siRNA design

To design siRNA targeting the gene of interest, a systematic bioinformatics approach was performed. First, the nucleotide sequence of the target gene was retrieved from the National Center for Biotechnology Information (NCBI) database. The coding sequence (CDS) was extracted in FASTA format. Next, potential siRNA candidates were designed using the RNAxs web tool (http://rna.tbi.univie.ac.at/cgi-bin/RNAxs/RNAxs.cgi). The extracted FASTA sequence was input into RNAxs with default design options, and the maximum number of siRNA was set to five. To ensure specificity, each proposed siRNA sequence (generally 19 to 21 nucleotides long) was subjected to a BLAST (Basic Local Alignment Search Tool) search (https://blast.ncbi.nlm.nih.gov/Blast.cgi) using the nucleotide BLAST (blastn) tool. Each siRNA was analyzed for sequence specificity by evaluating identity percentages, gaps, and alignment length. Only siRNA with high specificity to the target gene, minimal off-target interactions, and no significant matches with essential receptors or proteins were considered. Finally, selected siRNAs were synthesized and purchased through Eurogentec (see **Table S1**). The siRNA stock solutions were prepared in RNase-free water at 200 µM or 20 µM and stored at -20°C, as recommended by the provider. Each siRNA was tested for specificity and efficacy in targeting the gene of interest before selecting the most effective one for subsequent experiments.

### WRAP5:siRNA nanoparticle formulation

Nanoparticles were formulated in pure water supplemented by 5% glucose (Sigma-Aldrich) by mixing equal volumes of WRAP5 and siRNA at the corresponding molar ratio (WRAP5:siRNA = 20:1) at room temperature [42]. For PBN combining two siRNA (siYAP1+siNEG or siTAZ+siNEG or siYAP1+siTAZ), the two siRNA were pre-mixed according to siRNA stoichiometry 1:1 as described [44].

### Dynamic light scattering (DLS)

WRAP5:siRNA nanoparticles (WRAP5 = 10 µM, siRNA = 500 nM, R = 20) were evaluated with a Zetasizer NanoZS (Malvern) in terms of mean particle size (Z-average diameter) of the particle distribution and the polydispersity index (PdI). All results were obtained from three independent measurements (three runs for each measurement at 25°C) as described [42].

### Cell culture conditions

The experiments were conducted on human KIT-dependent IM-sensitive GIST cells. The GIST-T1 cell line (primary mutation in KIT exon 11 Δ560–578 in frame-deletion), from Cosmo Bio (Japan, Catalog No: PMC-GIST01C), was established from a metastatic Asian female human untreated GIST sample [49,50]. GIST-882 (KIT exon 13 K642E mutation) [51], and GIST-430 (KIT exon 11 Δ560-576 in-frame deletion) [52] were a gift from our collaborator, Dr. C. Serrano (Sarcoma Translational Research Laboratory, Vall d’Hebron Institute of Oncology [VHIO], Barcelona, Spain).

GIST-T1 cells were grown in Dulbecco’s Modified Eagle’s Medium (DMEM) (Corning, #10-013-CV) supplemented with 10% fetal bovine serum (FBS) (Sigma-Aldrich, #F7524) and 1% penicillin-streptomycin (Sigma-Aldrich, #P4333). GIST-430 and GIST-882 cells were cultured in Iscove’s Modified Dulbecco’s Medium (IMDM) (Sigma-Aldrich, #I3390) supplemented with 15% FBS, 1% penicillin-streptomycin, and 1% L-Glutamine (Thermo Fisher Scientific, #11524456). All the cells were passaged using Gibco™ Trypsin-EDTA (0.05%), phenol red (Thermo Fisher Scientific, #25300062). All the cells were maintained in a humidified incubator with 5% CO2 at 37°C. All cultures were tested monthly to be mycoplasma-free using the MycoAlert™ PLUS Mycoplasma Detection Kit (Lonza, #LT07-710). GIST-T1 cell lines were incubated with IM (STI571, Euromedex, France) at the concentrations indicated in the figure legends.

### GIST data set

Kaplan–Meier curves of progression-free survival stratified by *YAP1*, *TAZ*, *CYR61* or *CTGF* were obtained based on a clinically annotated gene expression data set (ATGsarc) of localized, untreated GIST (n = 60), quantified by microarray (ArrayExpress: E-MTAB-373) [53]. For each gene, patients were divided into high and low expression groups. The cut-off for stratifications was first set at the mean expression value, a standard approach in survival analysis to minimize bias from outliers (log-rank test). In parallel, box plot analyses were performed to compare the expression levels of these four genes between GIST patients with and without metastasis, providing additional insight into their potential association with disease aggressiveness.

### Cell cytotoxicity measurement

The cytotoxicity induced by the nanoparticles was evaluated using Cytotoxicity Detection Kit^Plus^ (LDH, Sigma-Aldrich) following the manufacturer’s instructions and as described before [42].

### Transfection experiments

For Western blot assays and RNA extraction, 300,000 GIST-T1 and 700,000 GIST-430 cells were seeded into 6-well plates (Sarstedt, #83.3920) 24 hours before the experiment. For GIST-882 cells, 500,000 cells were seeded 48 hours before the experiment into 6-well plates. For nanoparticle incubation, cells were treated with 800 µL of fresh, pre-warmed serum-free culture medium and 200 µL of nanoparticle solutions at the specified concentrations, or 200 µL of 5% glucose when required. After 1.5 h, 1,000 µL of the appropriate culture medium supplemented with 20% FBS for GIST-T1 (final FBS concentration = 10%) or 30% FBS for GIST-430 and GIST-882 (final FBS concentration = 15%) was added to each well, without removing the transfection reagents. The cells were incubated for an additional 24 hours or 48 hours before being lysed for RNA extraction or Western blot analysis, respectively.

### Western blotting

Transfected cells were treated as described before with some differences [44]. To standardize protein loading, all samples were diluted in RIPA buffer to match the lowest protein concentration. A 1:1 volume of 2X Laemmli Sample Buffer (Bio-Rad, #1610737) containing 5% β-mercaptoethanol was then added to each sample. Samples were heated at 95°C for 10 minutes, immediately placed on ice, and stored at -20°C until electrophoresis. Cell extracts were loaded with 5 to 10 µg of proteins. Electrophoresis and membrane transfer were achieved following [44]. Membranes were then blocked with TBS-0.5% or 0.2% Tween (Euromedex, #EU0660-B) containing 5% BSA (Sigma-Aldrich, #A2135). The 10X Tris-Buffered Saline (TBS) solution was prepared by dissolving 80 g NaCl, 2 g KCl, and 61 g Tris base (all from Sigma-Aldrich) in 800 mL of Milli-Q water, adjusting the pH to 8.0 with HCl, and completing the volume to 1 L with Milli-Q water. Membranes were subsequently incubated with the appropriate primary antibody in the same blocking buffer overnight at 4°C. After three washes with TBS-0.05% or 0.02%Tween , membranes were incubated for 1.5 h with the corresponding HRP-conjugated secondary antibody. Three additional washes were performed before chemiluminescent detection using luminol. The revelation was done following [44] and imaged using a Sapphire RGB & NIR Biomolecular Scanner (Azure Biosystems). VINCULIN, α-ACTININ, or β-ACTIN proteins were used as loading controls depending on the experiment. Primary and secondary antibodies were used according to the manufacturer’s recommendations. The details of the antibodies, including their source, catalog number, dilution, and application, are summarized in Supplemental Materials (see **Table S2**).

The signal intensities of the blots were quantified using Fidji ImageJ software as described in [44].

### Confocal microscopy

For confocal microscopy, 150,000 GIST-T1 cells were seeded onto 18 mm cover slips in 12-well plates (Sarstedt, #83.3921). 24 hours post-seeding, cells were incubated with 400 µL of fresh, pre-warmed serum-free DMEM and 100 µL of nanoparticle solutions at the specified concentrations. After 1.5 hours, 500 µL of DMEM supplemented with 20% FBS was added. After transfection, cells were fixed with 2% paraformaldehyde (PFA), and simultaneously, permeabilized and blocked using a solution of PBS containing 0.1% Tween-20/4% donkey serum (SIGMA, #D9663). Afterward, cells were incubated overnight with primary antibodies, followed by a 1-hour incubation with secondary antibodies. Nuclear staining was performed using Hoechst (1 mg/mL, Sigma, #14533), and membrane actin was labeled with Phalloidin Alexa Fluor™ Plus 647 (1:800, Thermo Fisher Scientific, #A22287). To visualize YAP1 and TAZ simultaneously, a mixed antibody staining approach was used. Finally, the cells were washed with D-PBS three times and mounted on glass slides with poly-(vinyl alcohol) Mowiol™ 4–88 glycerol Tris buffer (Biovalley). Confocal images were acquired using an inverted Zeiss LSM800 microscope with an Apo 63x/1.2 W DICIII objective. To minimize fluorophore crosstalk, sequential image acquisition was applied. For each biological replicate, 2 different arbitrary positions were selected for a z-stack mode acquisition (15 images with a z-stack interval of 0.22 µm). Acquired images were analyzed using Fiji (ImageJ) software. Sum of the fluorescence intensity for each z-stack projection was performed to obtain one image. The threshold of each sample was adjusted to quantify the mean fluorescence of the image. Mean fluorescence of YAP1 and TAZ was normalized by the mean fluorescence of the nucleus (Hoechst signal) and then adjusted to the WRAP5:siNEG condition.

### Cell Proliferation Assay by MTT

For MTT assay, 10,000 GIST-T1 cells were seeded into 96-well plates (Corning Falcon, #353072) 24 hours before the experiment. Cell proliferation was assessed using the CellTiter 96® Non-Radioactive Cell Proliferation Assay (Promega, #G4000), in which the luciferase-catalyzed luciferin/ATP reaction provides an indicator of cell number, based on the enzymatic conversion of MTT intoa colored formazan product. The manufacturer’s protocol was followed with slight modifications. The transfection with WRAP5:siRNA (1,200 nM:60 nM) and IM treatment (200 nM, Euromedex, #TO-I031) was performed in a total volume of 100 µL per well. Cells were incubated with 80 µL of serum-free DMEM and 10 µL of nanoparticle solutions. After 1.5 h, 10 µL of pure FBS was added, without removing the transfection reagents. All experiments were conducted in quintuplicate. At 24 h or 48 h post-transfection, 15 μL of the Dye Solution was added to each well, and the plate was incubated for 2 h. Subsequently, 100 µl of the Solubilization Solution/Stop Mix was added to each well, and the plate was incubated at room temperature for 1 h on a rocker to ensure a uniformly colored solution. Cell proliferation was measured using an Infinity M200 Pro microplate reader (Tecan Group Ltd) at 570 nm to quantify formazan formation and at 630 nm to minimize background interference from cell debris and nonspecific absorbance.

Cell proliferation was obtained using the following formulas:

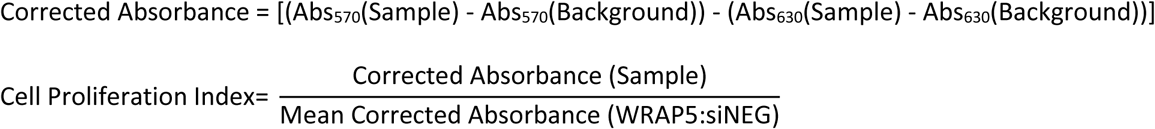

This approach normalizes the absorbance readings by removing background interference and comparing sample proliferation to the WRAP5:siNEG control. The background group was defined as the absorbance of wells containing medium without cells.

### Cell proliferation assay by cell counting

For cell proliferation analysis by cell counting, 100,000 GIST-T1 cells were seeded into 12-well plates 24 hours before the experiment. The transfection of WRAP5:siRNA (800 nM:40 nM) and IM treatment (50 nM) were performed in duplicate 24 hours after seeding. Transfection was performed using 400 µL of serum-free DMEM and 100 µL of nanoparticle solution. After 1.5 h, 500 µL of DMEM supplemented with 20% FBS was added. Measurements were taken at 0 h, 24 h, 48 h, and 72 h post-treatment following an established protocol. Before counting, viable cells were washed twice with DPBS 1X and incubated for 5 min with Trypsin-EDTA (0.05%) containing phenol red to detach the adherent cells. The resulting suspension was brought to a final volume of 1,000 µL by adding 800 µL of DPBS 1X. Proliferation was assessed using the CellDrop BFx automated counter (DeNovix, Wilmington, DE, USA) with the Brightfield program. A custom program optimized for GIST-T1 was used to ensure precise quantification. For each well, four independent counts were performed on 10 µL aliquots, and the average was used to determine the final cell concentration (cells/mL). To assess cell growth, counts at each time point were normalized to the mean count at t = 0 h.

### Wound healing assay

A wound healing assay was performed to assess the migratory capacity of GIST-T1 cells. To standardize the procedure, a Culture-Insert 4 Well (ibidi GmbH, #80469) was used in a 12-well plate. To minimize evaporation, 700 µL of medium was added to the area surrounding the insert. Next, 110 µL of a cell suspension containing 100,000 cells in complete DMEM was carefully added to each well of the insert. Transfection with WRAP5:siRNA (1,600 nM:80nM) was performed 24 h after seeding. The medium was gently removed, and 90 µL of serum-free medium was added, followed by 10 µL of WRAP5:siRNA nanoparticle solution. The cells were incubated for 1.5 hours at 37°C, after which 10 µL of pure serum was added to each well. After 24 hours, the insert was carefully removed from each well and filled with 1 mL of complete DMEM. If necessary, 1-2 washes with serum-free medium were performed to remove detached cells and debris.

Images of the wound area were captured immediately after scratching (time 0) and at 48 hours (72 hours post-transfection) using the EVOS® LX Core microscope system (#AMEX1000, Thermo Fisher Scientific), equipped with a 3.1 megapixel color camera (2048 x 1536 pixels, 1/2-inch sensor) and a resolution of 1024 x 768 pixels. The wound area was quantified using the Wound_healing_size_tool.ijm plugin in ImageJ, with adjusted parameters: Variance 10, Threshold 25, and Saturation 0.200. The percentage of wound closure was calculated by subtracting the area at t0 from the area at t48, and the results were normalized to the siNEG condition to assess cell migration.

### Reverse transcription and quantitative polymerase chain reaction (RT-qPCR)

Total RNA was extracted from cell cultures using the RNeasy® Mini Kit (Qiagen, # 74104), following the manufacturer’s protocol with some modifications. Specifically, cells were lysed in 230 µL of RLT buffer per well in a 6-well plate. The lysates were then processed according to the manufacturer’s instructions to obtain RNA samples. RNA quality and concentration were assessed using a NanoDrop™ One spectrophotometer (Thermo Scientific). The extracted RNA was then reverse transcribed into complementary DNA (cDNA) using 500 ng of RNA as the input material. The reverse transcription was carried out using the Verso cDNA Synthesis Kit (Thermo Scientific, #AB-1453/B), according to the manufacturer’s instructions. Following the synthesis, the cDNA was diluted in ultra pure RNase-free water to a final concentration of 2.5 ng/µL for use in subsequent qPCR analysis. Quantitative PCR (qPCR) was performed using the LightCycler® technology (Roche Diagnostics). PCR primers were designed using the LightCycler Probe Design 2.0 software for some targets, while others were sourced from the literature. Detailed primer sequences are provided in **Table S3**. To ensure primer specificity, each primer was verified through a BLAST search to confirm that there were no mismatches and that only the target of interest was amplified. Additionally, the efficiency of each primer was validated by performing a dilution series of cDNA ranging from 10 ng to 0.001 ng of cDNA per reaction. For each qPCR reaction, 5 ng of cDNA was added per well in a 96-well qPCR plate, in a final reaction volume of 10 µL. Reactions were carried out using the SYBR Green detection method (Roche Diagnostics, #04887352001), qPCR was conducted in technical triplicate, and standard curves were generated for each primer pair to assess amplification efficiency. Data were normalized to the reference gene *HPRT1*, and relative gene expression was calculated using the 2^−ΔΔCt method, which compares the Ct values (threshold cycle) of the target genes relative to the reference genes and the calibrator sample. All qPCR reactions were performed in a LightCycler® 480 System (Roche Diagnostics, #05015243001), and the results were analyzed and presented as mean gene expression levels relative to the reference genes.

### Statistical analysis

All statistical analyses were performed using GraphPad Prism (version 8.0.1). Data are presented as mean ± standard deviation (SD) or standard error of the mean (SEM) when appropriate, based on at least three independent experiments. Normality of data distribution was assessed using the Shapiro-Wilk test. When data did not follow a normal distribution, the Kruskal-Wallis test was used for multiple comparisons, as it is a non-parametric alternative to one-way ANOVA that does not assume normality, with Dunn’s correction applied for pairwise comparisons to either siNEG-treated cells or untreated cells. When residuals followed a Gaussian distribution and equal standard deviations were confirmed, a one-way ANOVA was performed for multiple comparisons, followed by Dunnett’s post-hoc test to compare each condition to either untreated cells or the siNEG-treated cells. In a case of two-group comparison, an unpaired two-tailed Student’s t-test was applied. A 95% confidence interval was applied to all tests, and statistical significance (p-value) was defined as follows: ns (not significant) >0.05, * <0.05, ** <0.01, *** <0.001, and **** <0.0001. In the figures, the number of biological replicates (N) and the number of measurements made within each biological replicate (n) are shown.

## RESULTS

### Role of YAP1 and TAZ in GIST prognosis

In our previous work, we identified LIX1 expression in GIST tissue microarrays in 79% of all GIST specimens (61/77) as well as in 79% of high-grade GIST (34/43) [17]. Moreover, *LIX1* was assessed as a negative prognostic factor in GIST, with high expression levels associated with shorter disease-free survival and increased metastatic risk. Importantly, LIX1 was suspected to regulate GIST cell plasticity through the modulation of YAP1 activity, suggesting a mechanistic link between these factors [17,54].

In the present study, we first evaluated whether the expression of *YAP1* and *TAZ* (*WWTR1*) was also associated with clinical outcome in GIST patients, as we previously did for *LIX1*. We analyzed a cohort of 60 patients from the ATGsarc microarray database [53]. The first analyzed data set comprised 28 GIST patients with high and 32 with low *TAZ* expression, and the second one contained 35 GIST patients with high and 25 with low *YAP1* expression (**Figure 1A**). Both blotted Kaplan-Meier curves revealed that a tendentially high expression of *YAP1* (*p* = 0.054) or a significantly high expression of *TAZ* (*p* = 0.0021) could be associated with shorter progression-free survival, indicating a potential role in tumor progression.

**Figure 1:**
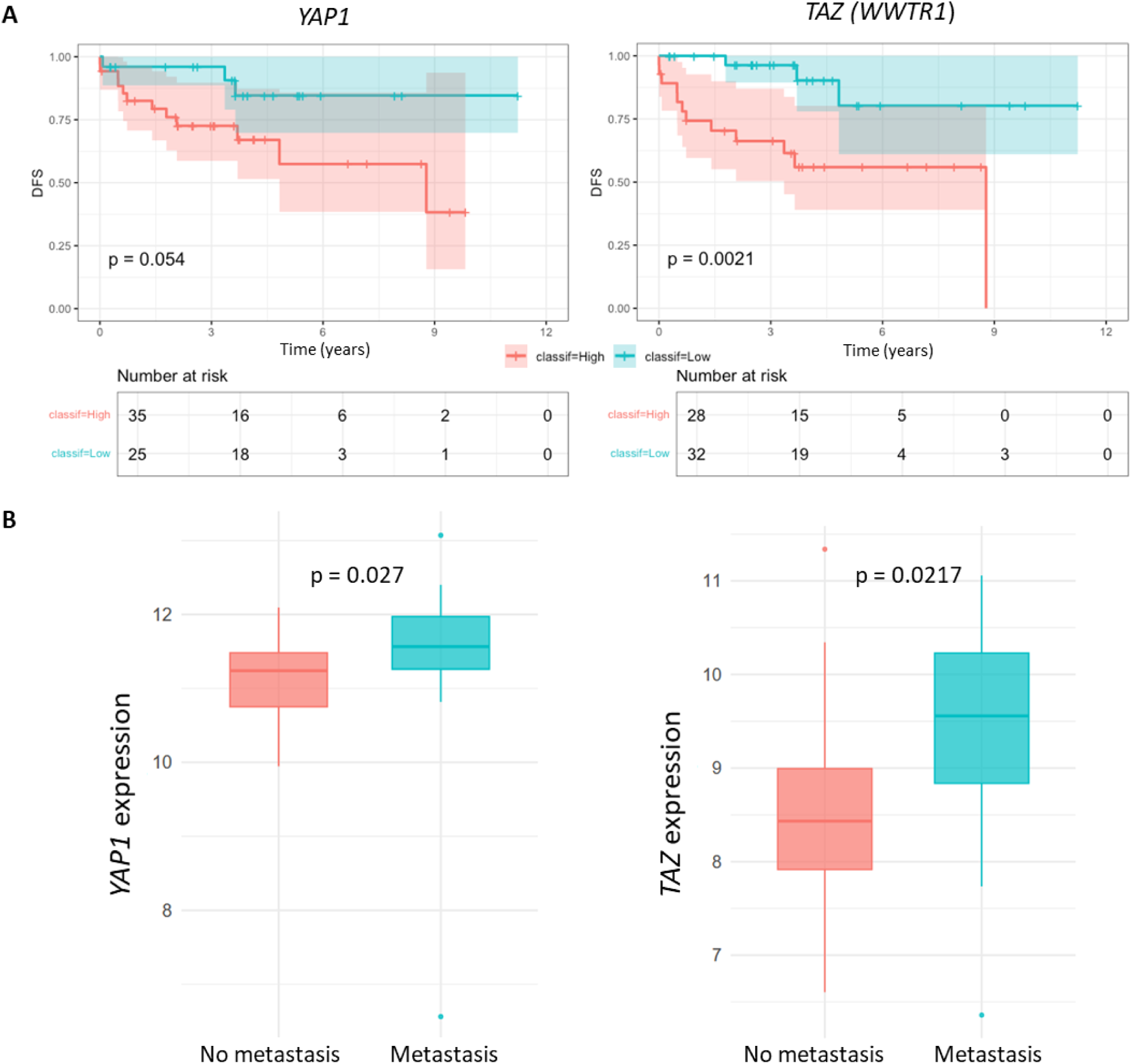
Correlation of *YAP1* and *TAZ* expression with the prognosis of GIST patients. (**A**) Kaplan-Meier curves of disease-free survival (DFS) from 60 total GIST patients in the ATGsarc microarray database, stratified by *YAP1* or *TAZ* (*WWTR1*) expression levels. Group 1 (red): patients with high gene expression (number initial of patients = 35) ; Group 2 (blue): patients with low gene expression (number initial of patients = 25). Number at risk = number of patients at risk over time. They are resumed in the table below the curves. (**B**) Comparison between *YAP1* or *TAZ* transcript levels and the presence of metastasis. n = 45 (red) and n = 15 (blue) in no metastatic and metastatic groups, respectively. p-values from the log-rank test are indicated.

In parallel, the *YAP1* and *TAZ* high transcript levels were correlated to the presence or absence of metastases. Both analyzed datasets comprised 15 GIST patients with and 45 without metastases (**Figure 1B**). Both the boxplot indicated a significant correlation of *YAP1* expression (*p* = 0.027) or *TAZ* expression (p = 0.0217) and metastasis formation , linking both genes to an aggressive tumor behavior.

Collectively, these results supported a model in which *TAZ* and, to a lesser extent, *YAP1* contributed to GIST progression. These observations led us to functionally assess the individual contributions of *YAP1* and *TAZ* in GIST cells, using a gene-specific silencing approach *in vitro*.

### Specific silencing of YAP1 or TAZ using WRAP5:siRNA nanoparticles in GIST-T1 cells

To selectively investigate the individual and combined roles of YAP1 and TAZ in KIT-dependent IM-sensitive GIST-T1 cells, we designed for each gene different siRNA using the RNAxs web tool. Each siRNA was blasted to ensure the specificity of the sequence for its target. Two siRNA with the highest score, each targeting two different complementary region of the targeted mRNA, were selected for evaluation (**Table S1**).

For the cellular internalization of the siRNA, we used the WRAP5-based nanoparticle delivery system previously validated for efficient and low-toxicity gene silencing across diverse cancerous and non-cancerous cellular models [42,43,46]. First, we used dynamic light scattering (DLS) to validate WRAP5:siRNA nanoparticle integrity, showing an average diameter ranging from 90 to 120 nm, with a polydispersity index (PdI) of around 0.3 (**Figure S1**). These relatively homogeneous populations of nanoparticles were consistent with previous reports [42,44] and were, therefore, suitable for gene silencing experiments.

YAP1 and TAZ silencing were evaluated in GIST-T1 cells using WRAP5:siYAP1 and WRAP5:siTAZ in a dose-dependent manner (20 nM to 80 nM) by Western blot (**Figures 2A** and **2B**). This analysis was performed compared to non-treated GIST-T1 cells as well as to those incubated with WRAP5 nanoparticles encapsulating a siRNA without any cellular target (siNEG) to ensure that the nanoparticles have no influence on cell physiology.

**Figure 2:**
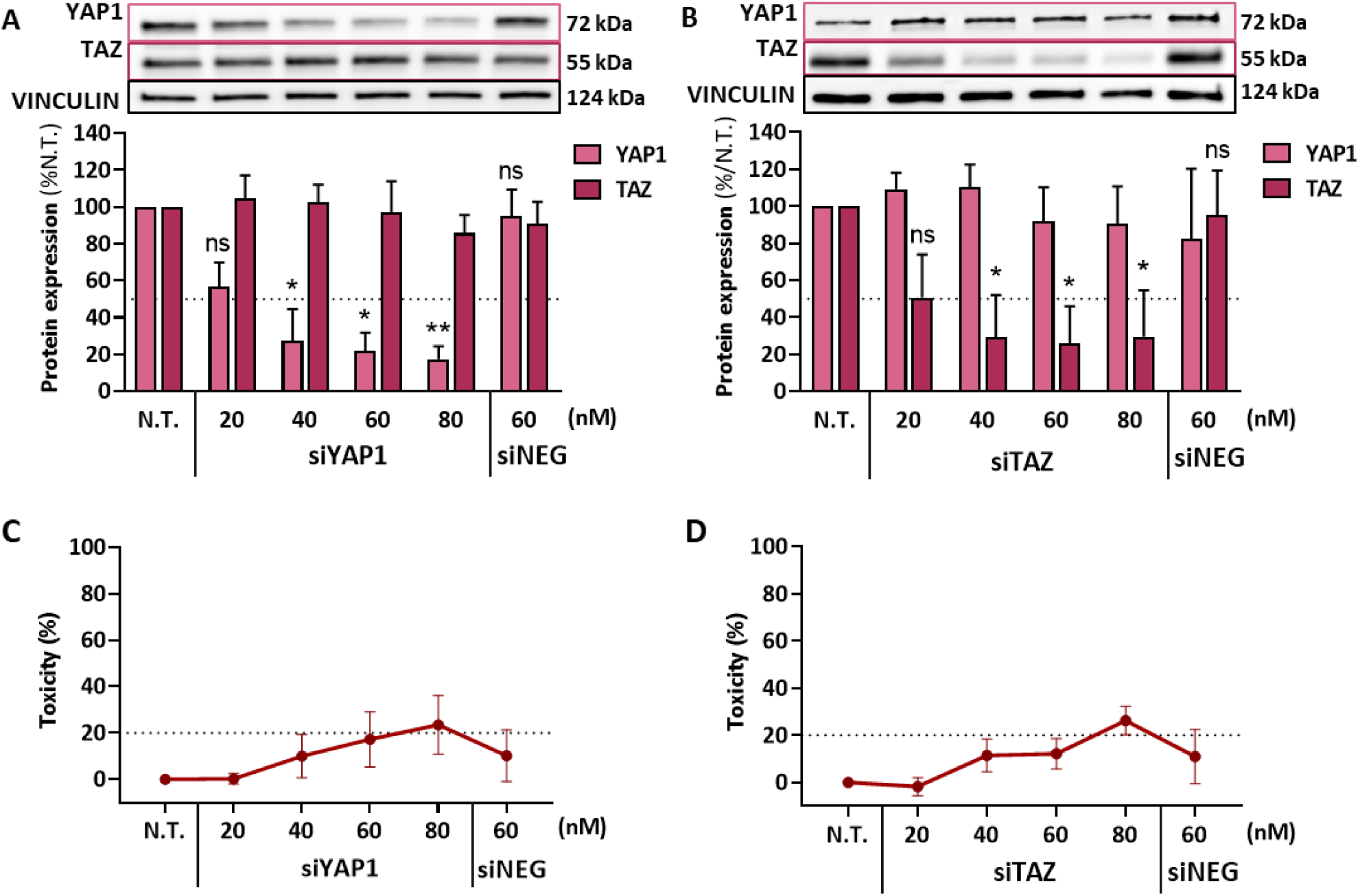
Evaluation of YAP1 or TAZ silencing in GIST-T1 cells with WRAP5:siRNA nanoparticles. WRAP5:siRNA nanoparticles delivering siYAP1 (**A**) or siTAZ (**B**) induced a dose-dependent inhibition of YAP1 or TAZ protein expression in GIST-T1 cells after 48 h of incubation, as shown by Western blot quantification. No significant toxicity was observed for siYAP1 (**C**) or siTAZ (**D**) within the tested concentration range (20 – 60 nM), as assessed by LDH assay. A slight toxicity was observed for 80 nM. Controls included untreated (N.T.) and siNEG-treated cells. Data are presented as mean ± SD from N=4 experiments. Statistical analysis was performed using Kruskal-Wallis followed by Dunn’s multiple comparisons test *versus* N.T. (**A**, **B**); ns >0.05, * <0.05, ** <0.01. Analysis of TAZ expression following siYAP1 treatment and YAP1 expression following siTAZ treatment showed no significant differences compared to N.T. (data not annotated on the graph for clarity).

Concerning the YAP1 and TAZ silencing, we observed, in both cases, a specific knockdown of YAP1 with the WRAP5:siYAP1 nanoparticles without any effect on the TAZ protein expression (**Figure 2A)**. The same specificity has been shown for WRAP5:siTAZ nanoparticles on the TAZ and YAP1 expression (**Figure 2B)**. This knockdown specificity was further confirmed with additional controls: (1) alternative siRNA sequences targeting another region of the *YAP1* and *TAZ* mRNA showed no cross-silencing between YAP1 and TAZ, depending on the siRNA used (**Figures S2A, S2B**), and (2) WRAP5:siNEG nanoparticles did not induce any YAP1 or TAZ silencing compared to the untreated GIST-T1 cells **Figures 2A, 2B, S2A, S2B**).

For both proteins, YAP1 and TAZ, the maximal silencing efficacy reached a plateau of approximately 70% at a siRNA concentration of 60 nM, depending on the used siRNA (**Figures 2A, 2B, S2A, S2B**). To confirm that this range of silencing was correct for both proteins, we encapsulated a siRNA (siYAP1TAZ) [55], that silence both proteins (**Figure S3**). Using this siRNA, a silencing of around 70% for both proteins was revealed, confirming the specificity and efficacy of our designed siRNA.

To ensure that the nanoparticles did not interfere with the cell viability, we performed for all analyzed conditions (**Figures 2A, 2B, S2A, S2B**) cytotoxicity assays (LDH) before cell lysis for Western blot. Indeed, we revealed minimal cytotoxicity below 20% for siRNA concentrations up to 60 nM and only some slight effects (20%-30%) using the highest siRNA concentration of 80 nM (**Figures 2C, 2D, S2C, S2D**). These thresholds for cytotoxicity were in the acceptable range as described in the standard ISO 10993-5 used to evaluate the *in vitro* cytotoxicity of medical devices [56].

Finally, we compared YAP1 and TAZ expression levels depending on different percentages of GIST-T1 cell confluence (30%, 50%, 80%, and 100%) to ensure that, for both, protein quantity will be the same under different transfection protocols used in the next experiments. Western blots analyses confirmed that, at working cell confluence ranges between 50% and 100%, no significant difference between YAP1 or TAZ expressions were observed (**Figure S4**). This finding confirmed that GIST-T1 cell confluence and potent resulting contact inhibition did not affect the protein quantity of YAP1 and TAZ [57,58].

### Simultaneous silencing of YAP1 and TAZ in GIST-T1 cells

Knowing that the WRAP5:siYAP1 and the WRAP5:siTAZ nanoparticles could perform specific YAP1 or TAZ silencing (at 40 nM siRNA), we wanted to combine both siRNA to further evaluate if a phenotypic synergistic effects could be observed. The delivery of two or three siRNA simultaneously by the WRAP5 nanoparticles was previously described in glioblastoma U87 cells, but also in GIST-T1 cells [44]. Furthermore, as the WRAP5:siNEG nanoparticles had no effect on cell physiology and cytotoxicity, we decided to use this condition for the result normalization for all further assays.

The siRNA cocktail with an equimolar ratio (1:1) of siYAP1 and siTAZ (WRAP5:siYAP1+siTAZ) was formulated with a constant siRNA concentration (40 nM) across conditions, meaning that [20 nM siYAP1+20 nM siTAZ] were used for the (WRAP5:siYAP1+siTAZ) condition (**Figures 3A, 3B**). Western blot analyses showed a significant YAP1 (*p* <0.01) and TAZ (*p* <0.05) silencing, respectively, of around 50%, compared to WRAP5:siNEG, which was in coherence with the results shown in **Figures 2A** and **2B**. The 50% silencing was also observed when one the two loaded siRNA was replaced by the siNEG, showing that there was probably no cross-talk between both silenced proteins.

**Figure 3:**
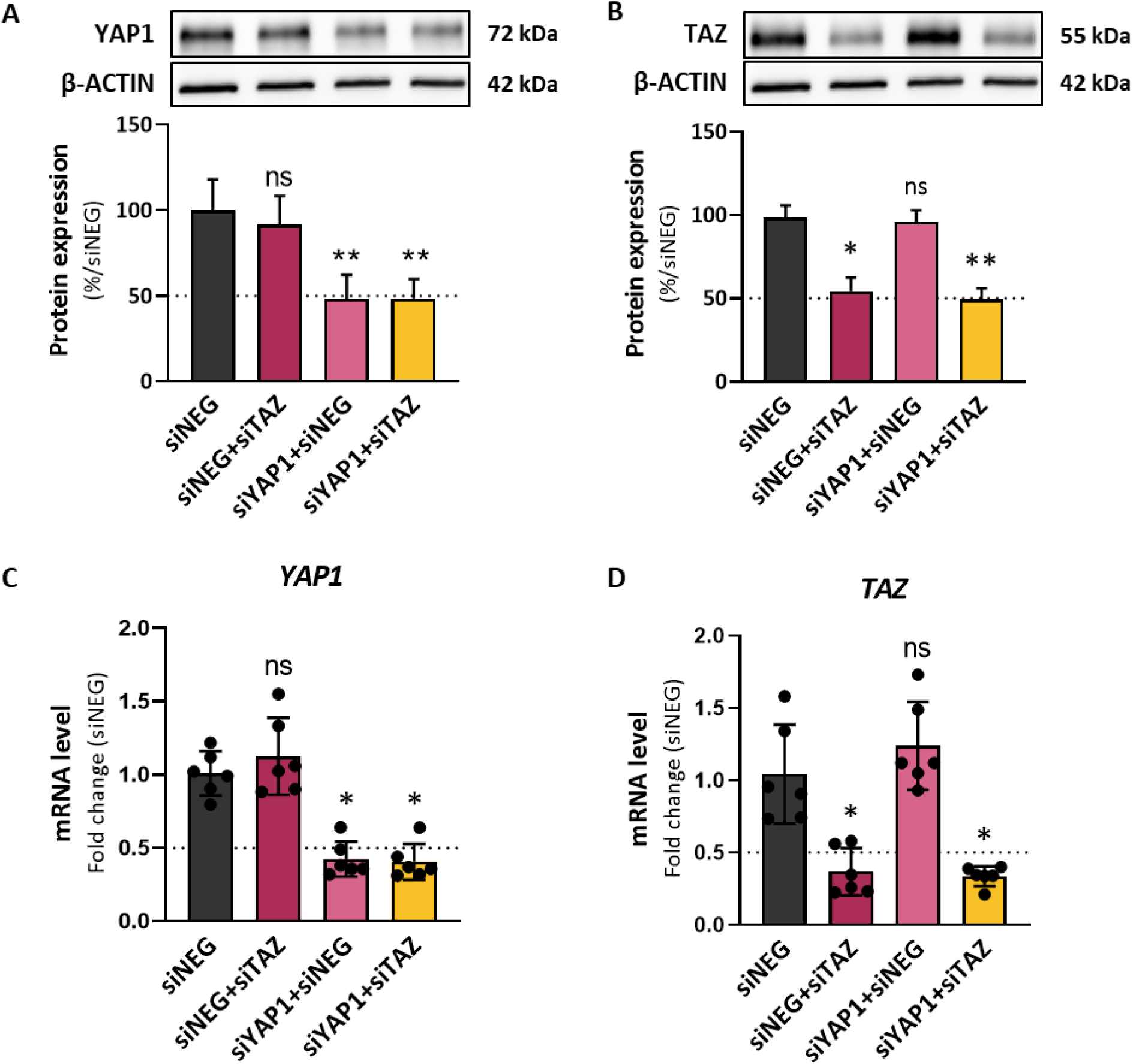
Specific targeting of YAP1 and TAZ, individually or simultaneously, using WRAP5:siRNA cocktails. WRAP5:siRNA nanoparticles delivering an equimolar cocktail of siRNAs targeting YAP1 and/or TAZ specifically inhibited YAP1 and TAZ protein expression (**A, B**) and mRNA levels (**C, D**) in GIST-T1 cells, as assessed by Western blot quantification and qPCR analysis, respectively. Conditions: WRAP5:siRNA (MR 20 with [siRNA] = 40 nM), siNEG-treated cells were used as a control condition. Data were presented as mean ± SD for N=6 individual experiments. Statistical analyses were performed using Kruskal-Wallis followed by Dunn’s multiple comparisons test *versus* siNEG; ns >0.05, * <0.05, ** <0.01.

Analyses by qPCR confirmed the robust silencing (up to 70%) of *YAP1* (*p* <0.05) and *TAZ* (*p* <0.05) mRNA when the WRAP5 nanoparticle encapsulated both siRNA (siYAP1+siTAZ) compared to WRAP5:siNEG (**Figures 3C, 3D**). Here again, comparable mRNA silencing efficiency was observed for siRNA delivered alone or in combination, allowing us to rule out any cross-talk due to simultaneous knockdown of YAP1 and TAZ.

To further confirm the specificity and efficiency of YAP1 and TAZ silencing at the cellular level, we performed immunofluorescence staining of both proteins in GIST-T1 cells treated with WRAP5 nanoparticles loaded with siNEG, siYAP1, siTAZ, or siYAP1+siTAZ (**Figure 4A**). Cells were co-stained for YAP1 and TAZ, and fluorescence intensity was quantified across individual cells. A significant decrease in TAZ signal (-45%, *p* <0.01) was observed in cells treated with siTAZ (**Figure 4B**), while YAP1 signal was significantly reduced (- 40%, *p* <0.01) in cells treated with siYAP1 (**Figure 4C**) compared to the siNEG condition. Notably, no nuclear relocalization or compensatory accumulation of one protein was observed when the other was silenced. This absence of compensatory nuclear recruitment suggests that YAP1 and TAZ do not substitute for each other in GIST-T1 cells under these conditions. However, it seems that the fluorescence signal of TAZ is significant lower (-64%, *p* <0.0001), when both siRNA (siYAP1+siTAZ) were used, in contrast to the YAP1 fluorescence signal which was constant (-40%, *p* <0.01; equal to siYAP1 alone) (**Figures 4B, 4C**).

**Figure 4:**
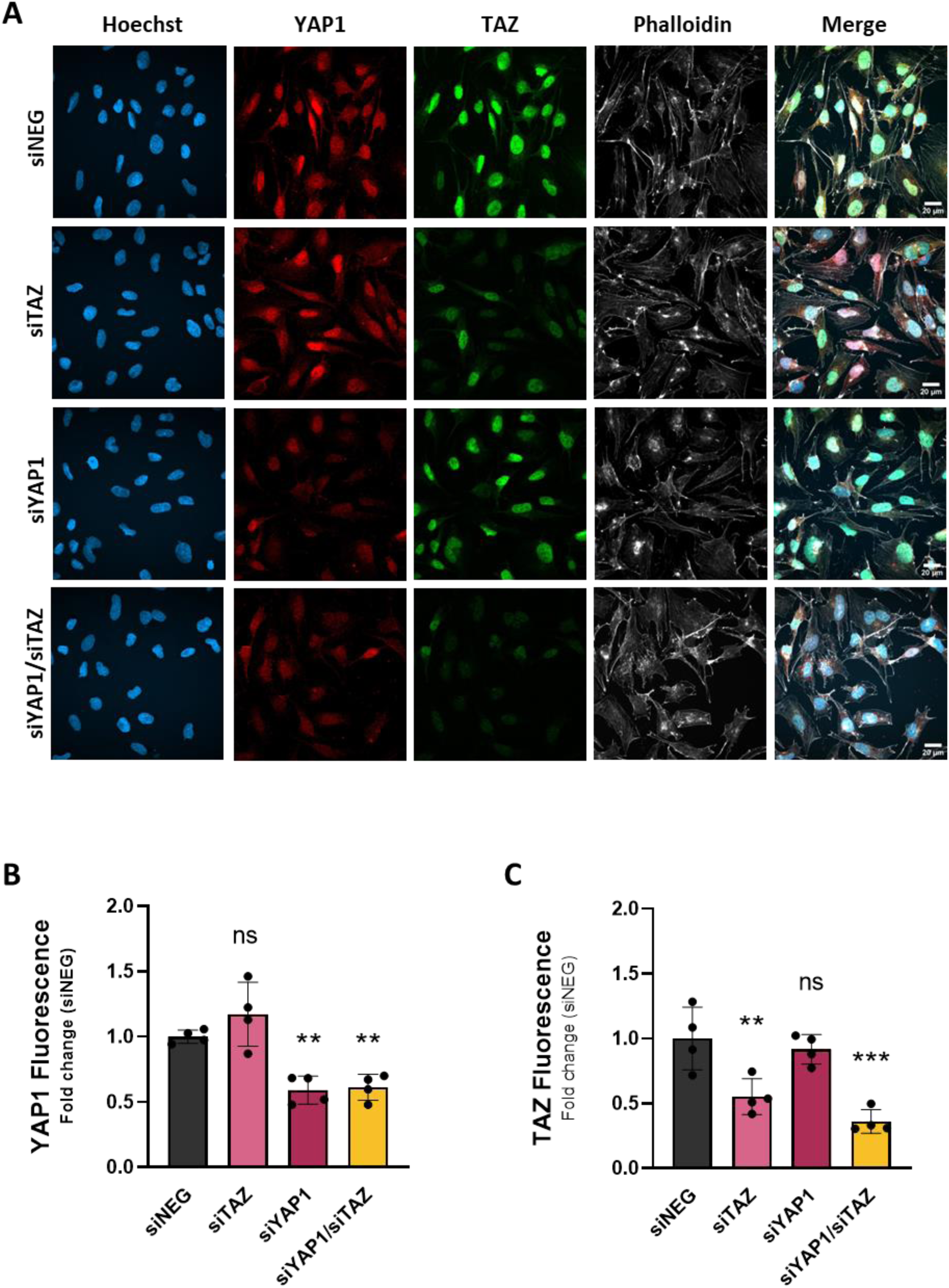
Immunofluorescence analysis of YAP1 and TAZ expression following siRNA-mediated silencing in GIST-T1 cells. GIST-T1 cells were treated with WRAP5 nanoparticles loaded with 40 nM of siNEG (control), 20 nM of siYAP1 or siTAZ, or a combination of siYAP1+siTAZ (20 nM each). Immunofluorescence staining was performed to detect endogenous YAP1 (red) and TAZ (green), with nuclei counterstained using Hoechst (blue) and Phalloidin (gray). Representative images are shown for each condition (**A**). Scale bars = 20 μm. Quantification of fluorescence intensity by Image J were performed for YAP1 (**B**) and for TAZ (**C**) in the four conditions. Data were represented as mean ± SD for N = 2 with n=2. Statistical analyses were performed using Kruskal-Wallis followed by Dunn’s multiple comparisons test *versus* siNEG.

These results established WRAP5:siRNA nanoparticles as a reliable and non-toxic tool for selective and combinatorial gene silencing of YAP1 and TAZ in GIST cells, enabling us to dissect the role of both proteins individually or together in GIST biology.

### Selective effect of TAZ silencing on GIST-T1 cell migration and proliferation

After proving the effectiveness of our WRAP5:siRNA nanoparticles in silencing YAP1 and TAZ individually or together in GIST-T1 cells, we next investigated the functional consequences of their knockdown on the cellular mechanisms, such as cell migration and proliferation.

To assess cell migration, we performed a wound healing assay on confluent GIST-T1 cells seeded on a culture insert to reproduce a homogeneous scratch (wound). 72 hours after transfection with siRNA-loaded WRAP5 nanoparticles, we observed that TAZ silencing resulted in a substantial decrease (-24%) in wound closure compared to the siNEG control condition (*p* <0.01), suggesting an impaired migratory capacity (**Figure 5A**). With regard to YAP1 silencing, it is interesting to note that no significant effects (p = ns) has been observed, despite previous reports of its involvement in migration phenomena in other cancers [59,60]. Co-silencing of YAP1 and TAZ using WRAP5:(siYAP1+siTAZ) resulted in cell migration reduction compared to the siNEG control (-31%, *p* <0.001) (**Figure 5**). This slightly higher wound healing revealed that the co-silencing may have an additive effect compared to TAZ silencing alone, as observed previously in confocal microscopy (**Figure 4**).

**Figure 5:**
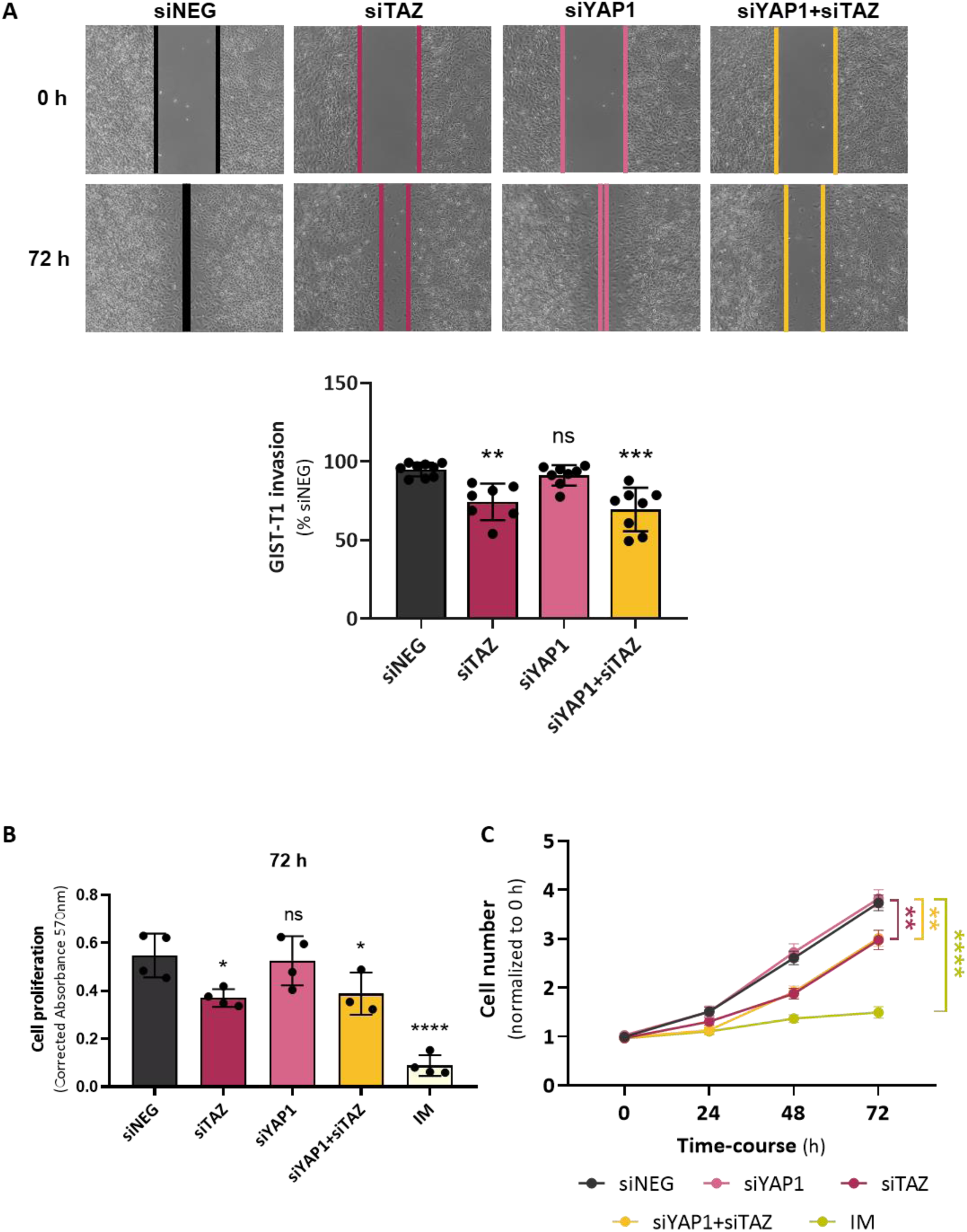
TAZ silencing reduced GIST-T1 cell migration and proliferation. (**A**) WRAP5:siRNA nanoparticles targeting TAZ, YAP1 or both (siYAP1+siTAZ) were analyzed in wound healing assay 72 h post-transfection in GIST-T1 cells. The graph shows the percentage of wound closure relative to the initial scratch and normalized to the siNEG condition (N=8 independent experiments). (**B**) WRAP5:siRNA nanoparticles targeting TAZ, YAP1 or both (siYAP1+siTAZ) were analyzed using the CellTiter 96® Non-Radioactive Cell Proliferation Assay 72 h post-transfection in GIST-T1 cells (N=4 independent experiments with n=5 replicates). (**C**) WRAP5:siRNA nanoparticles targeting TAZ, YAP1 or both (siYAP1+siTAZ) were analyzed by cell counting over a 72 h time course post-transfection (N=5 independent experiments with n=2 replicates). Abbreviation : IM = Imatinib. Conditions: WRAP5:siRNA (MR 20) with siRNA concentrations as indicated in the Material & Methods section, siNEG was used as the control condition, IM = 200 nM. Data are presented as mean ± SD (**A**, **B**) or mean ± SEM (**C**). Statistical analyses were performed using Kruskal-Wallis followed by Dunn’s post-hoc test *versus* siNEG (**A**); one-way ANOVA followed by Holm-Sidak’s post-hoc test *versus* siNEG (**B**,**C**). ns >0.05, * <0.05, ** <0.01, *** <0.001, and **** <0.0001.

We next evaluated cell proliferation firstly using the MTT assay, an approach that measured metabolic cell activity with the same incubation conditions as used for the wound healing assay (**Figure 5B**). Evaluation of the assay 48 hours post-transfection showed no statistical differences between the WRAP5-based conditions (**Figure S5**). Only GIST-T1 cells incubated with IM (200 nM used as control) revealed a significant reduction in cell proliferation (-85%, *p* <0.0001). In contrast, when the assay was evaluated 72 hours post-transfection, a marked decline in proliferation rate was observed for TAZ silencing (-30%, *p* <0.05) in comparison to the control condition siNEG or siYAP1. Interestingly, we observed an equivalent effet on proliferation when GIST-T1 cells were co-treated with (siYAP1+siTAZ) (-20%, *p* <0.05).

Similar results were observed when performing a direct cell counting over time (**Figure 5C**). Following transfection with WRAP5-based nanoparticles, YAP1 silencing alone did not result in a statistically significant reduction in cell proliferation. However, a reduction in the number of cells was observed 48 hours or 72 hours post-transfection when TAZ was silenced alone or in combination with YAP1 (around -21% at 72 h, *p* <0.01). No potential additive effect of the applied siYAP1+siTAZ combination could be detected in both cell proliferation assays.

Together, these findings revealed that TAZ, more than YAP1, could drive GIST-T1 cell proliferation and migration, and that its silencing had a stronger inhibitory effect on these tumorigenic properties.

### Impact of TAZ silencing on Hippo pathway effectors in GIST-T1 cells

Following the observed phenotypic effects of TAZ silencing on KIT-dependent IM-sensitive GIST-T1 cell migration and proliferation, we aimed to explore the molecular mechanisms underlying its function. Following activation by phosphorylation, YAP1 and TAZ translocate into the nucleus, where they bind to the TEAD1-4 proteins, thereby facilitating the expression of downstream target genes (*CTGF, CYR61, AREG, MYC, MCL-1, BIRC5, AXL*, etc.) implicated in cell migration, proliferation, invasion and resistance to apoptosis [23–25].

As we observed an effect of TAZ on cell migration and proliferation, we first examined the expression of CYR61 and CTGF. Western blot analyses on GIST-T1 cells incubated with the WRAP5 nanoparticles encapsulating the siYAP1 alone induced no downstream effect on CYR61 and CTGF expression (*p* = ns). Conversely, WRAP5:siTAZ nanoparticles revealed a significant downregulation of CYR61 protein (-65%, *p* <0.001) and of CTGF (-55%, *p* <0.01) compared to the control condition (WRAP5:siNEG) (**Figures 6A, 6B**). This effect on CYR61 and CTGF protein expression was also observed when siTAZ was used in combination with siYAP1. All these results were confirmed by qPCR analyses that revealed a significant downregulation of *CYR61* and *CTGF* genes on mRNA levels upon TAZ knockdown, either alone or in combination with YAP1 (>80%, *p* <0.0001 for both respectively) compared to WRAP5:siNEG. In contrast, YAP1 silencing alone had no measurable effect on *CYR61* mRNA level, but, curiously, we observed an increase of *CTGF* mRNA upon siYAP1 silencing (**Figure 6D**). These indicated that TAZ, rather than YAP1, was the main driver of Hippo pathway output in GIST-T1 cells.

**Figure 6:**
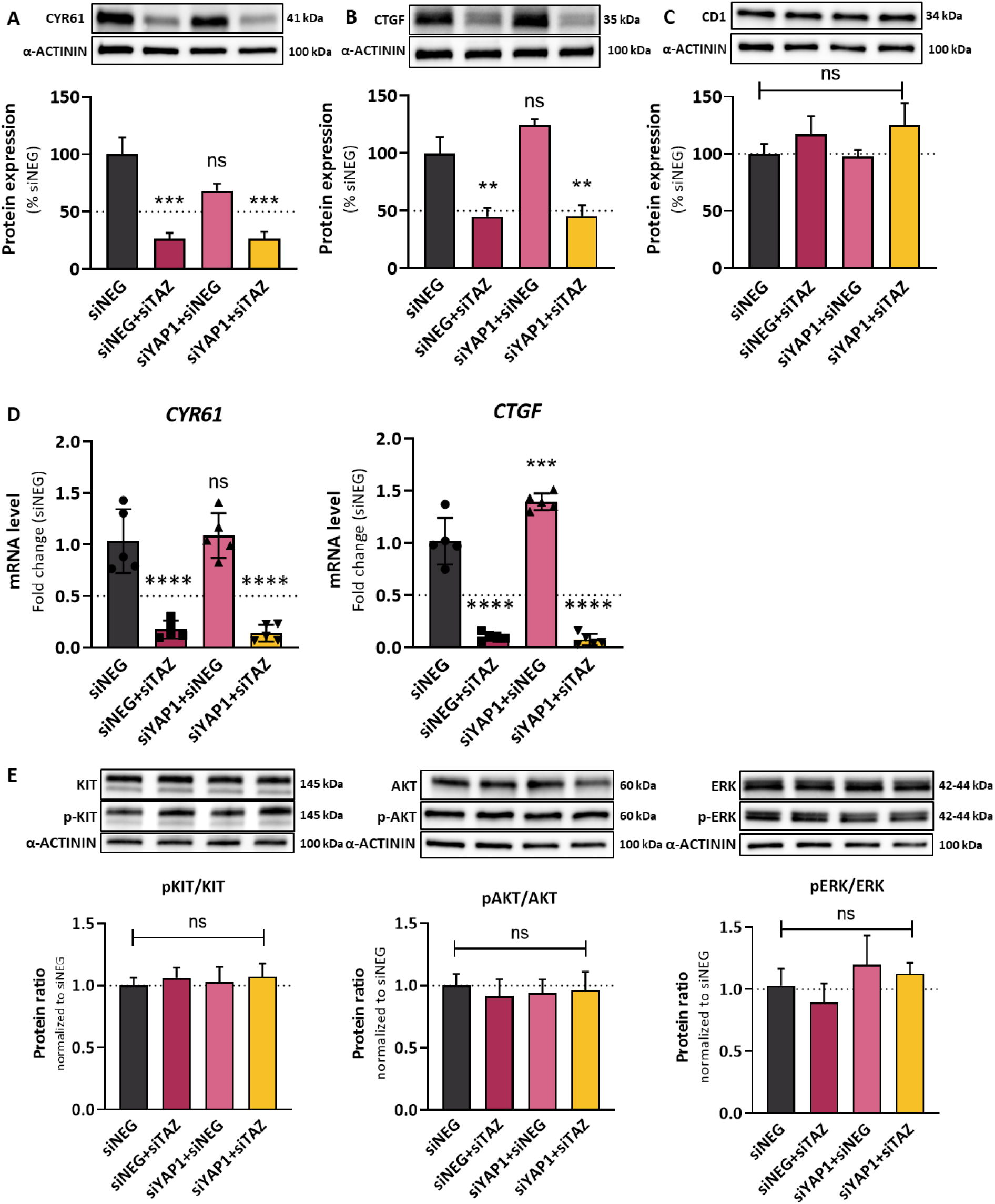
Evaluating TAZ silencing on different signaling pathways in GIST-T1 cells. Effect of WRAP5:siRNA nanoparticles targeting TAZ, YAP1 or both (siYAP1+siTAZ) on CYR61 (**A**), CTGF (**B**) and CD1 (**C**) protein expression in GIST-T1 cells were analyzed by Western blot analysis 48 h post transfection. (**D**) Effect of WRAP5:siRNA nanoparticles targeting TAZ, YAP1 or both (siYAP1+siTAZ) on CYR61 and CTGF mRNA levels in GIST-T1 cells were analyzed by qPCR analysis 24 h post transfection. (**E**) Effect of WRAP5:siRNA nanoparticles targeting TAZ, YAP1 or both (siYAP1+siTAZ) on of KIT signaling components (KIT, pKIT, AKT, pAKT, ERK, pERK) in GIST-T1 cells were analyzed by Western blot analysis 48 h post transfection. Conditions: WRAP5:siRNA (MR 20 with [siRNA] = 40 nM), siNEG was used as the control condition. Data are presented as mean ± SD with N = 4 - 6 individual experiments. Statistical analyses were performed using one-way ANOVA followed by Dunnett’s post-hoc test *versus* siNEG (**A**,**B,C,D**); Kruskal-Wallis followed by Dunn’s multiple comparisons test *versus* siNEG (**E**) ns >0.05, ** <0.01, *** <0.001, ****<0.0001.

We also analyzed Cyclin D1 (CD1) expression to determine whether this key cell cycle regulator was modulated by YAP1 or TAZ in our model. Surprisingly, CD1 expression remained unchanged following TAZ or YAP1 silencing (*p* = ns for all conditions *versus* siNEG), suggesting that its regulation in GIST-T1 cells was independent of these factors under the tested conditions (**Figure 6C**).

Finally, to investigate potential connection between the Hippo and KIT signaling pathways, we assessed ratios of total and phosphorylated levels of KIT, AKT, and ERK, mimicking the activation of MAPK and PI3K/mTOR pathways. They represent the key effectors of KIT oncogenic signaling [24,33]. No significant changes were observed following YAP1 or TAZ knockdown in GIST-T1 cells (**Figure 6E**), indicating that the anti-tumorigenic effects of TAZ silencing were independent of direct modulation of KIT or its canonical downstream cascades.

In conclusion, these results showed that TAZ promoted GIST-T1 proliferation and migration primarily through activation of specific Hippo pathway targets, rather than by influencing CD1 and the KIT signaling axis. We thus hypothesized that CYR61 and CTGF were specific downstream effectors of TAZ in GIST-T1 cells.

### WRAP5:siRNA as an effective tool for YAP1/TAZ silencing other GIST cells

As the GIST pathology was characterized by different mutations resulting in KIT-dependent IM-sensitive GIST, we decided to quantify YAP1 and TAZ expression in GIST-430 and GIST-882 cells [51,52]. These two cell lines harbors different *KIT* mutations than the primary mutation in GIST-T1 cell (*KIT* exon 11 Δ560– 578 in frame-deletion versus respectively KIT exon 11 Δ560-576 in-frame deletion and KIT exon 13 K642E mutation). These cells displayed distinct basal expression profiles of YAP1, and TAZ compared to GIST-T1 cells, as confirmed by Western blot analyses (**Figure S6**). Notably, TAZ expression was markedly lower in GIST-882 cells, while both YAP1 and TAZ were expressed at higher levels in GIST-430, underscoring the molecular heterogeneity that characterized GIST tumors [61]. Importantly, total and phosphorylated KIT levels appeared comparable across the three cell lines, suggesting that differences in YAP1 and TAZ expression and downstream target regulation were not attributable to variations in KIT signaling (**Figure S6**). However, given the differing morphologies (**Figure S7**) and proliferation rates of the three GIST cell lines, the protocol was adapted to each cell line in terms of cell density and siRNA concentrations (see Material & Methods section for details).

Using WRAP5:siRNA nanoparticles previously used, we evaluated YAP1 and TAZ silencing depending on the applied siRNA (siYAP1, siTAZ, siYAP1+siTAZ, siNEG) in GIST-430 and GIST-882 cells. In both cell lines, YAP1 and TAZ knockdown were efficient when the corresponding siRNA was used compared to the siNEG (between 40% and 70%, depending on the condition used) (**Figures 7A, 7B**). Surprisingly, for the WRAP5 nanoparticles delivering the equimolar siRNA cocktail of siYAP1 and siTAZ, we observed a slightly higher protein knockdown as with the single siRNA, which was not observed previously on GIST-T1 cells. However, when applying the one-way ANOVA statistical test followed by Dunnett’s post-hoc test with multiple comparisons versus siTAZ or (siYAP1+siTAZ), no significant difference was observed between these groups (data not shown).

**Figure 7:**
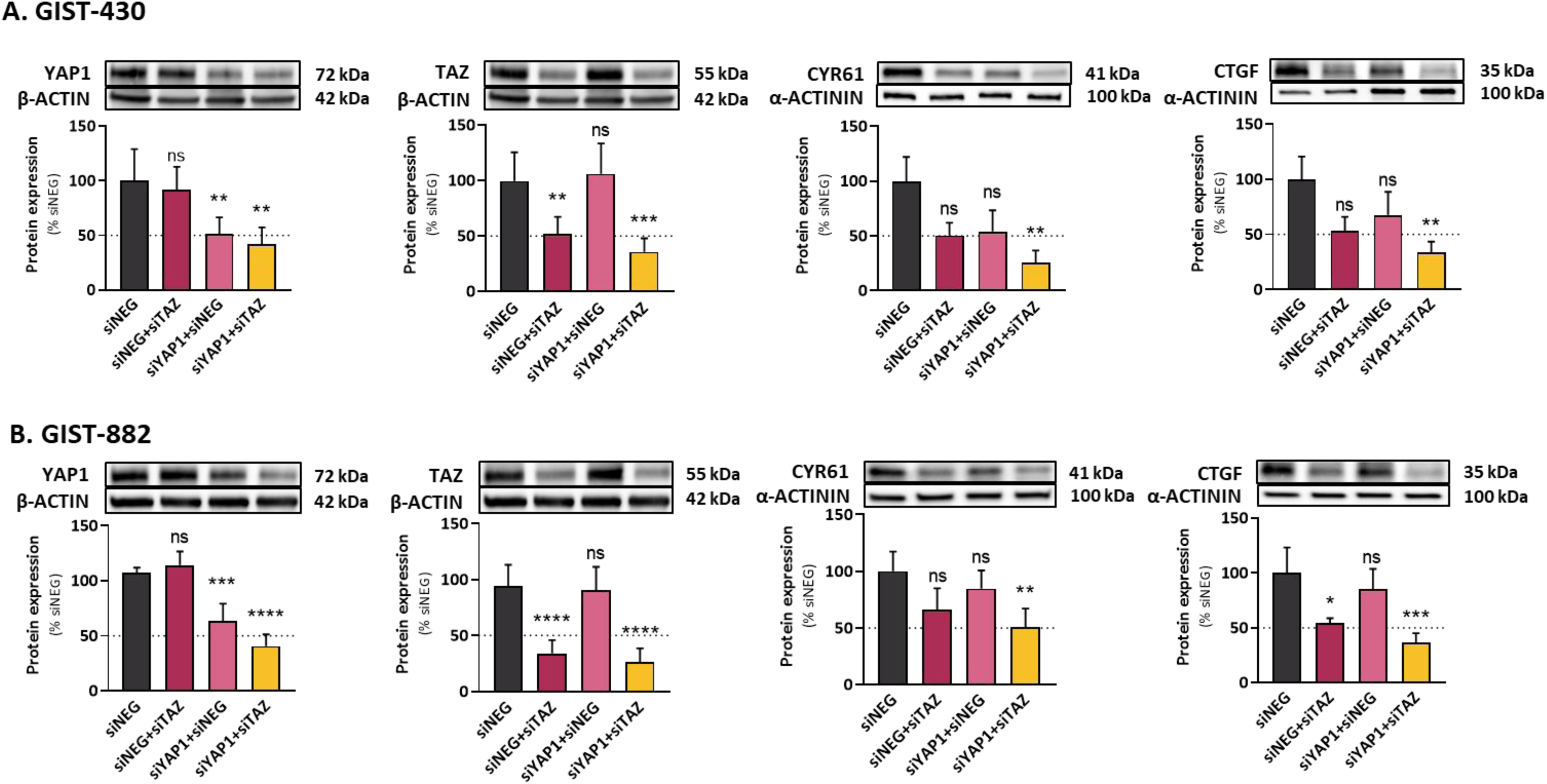
WRAP5:siRNA nanoparticles enabled selective silencing of YAP1 and TAZ across GIST cell lines, with conserved downstream regulation of CYR61 and CTGF. Effect of WRAP5:siRNA nanoparticles targeting TAZ, YAP1 or both (siYAP1+siTAZ) on YAP1 and TAZ or CYR61 and CTGF protein expression in GIST-430 (**A**) and GIST-882 (**B**) cells were analyzed by Western blot 48 h post transfection Conditions: WRAP5:siRNA (MR 20 with [siRNA] = 80 nM), siNEG was used as control. Data are presented as mean ± SD for N=5 individual experiments. Statistical analyses were performed using one-way ANOVA followed by Dunnett’s post-hoc test *versus* siNEG (**B,C**); Kruskal-Wallis followed by Dunn’s multiple comparisons *versus* siNEG (**D,E**); ns >0.05, * <0.05, ** <0.01, *** <0.001, **** <0.0001.

Then, we also analyzed YAP1/TAZ silencing on downstream protein expression of CYR61 and CTGF in GIST-430 and GIST-882 cells as previously done for GIST-T1 cells. In details, in GIST-430 cells, YAP1 or TAZ silencing alone led to a non-significant reduction in CYR61 and CTGF even if there was a trend for both proteins (*p* = ns *versus* WRAP5:siNEG), while their combination (siYAP1+siTAZ) significantly decreased both targets (50%, *p* <0.01 for CYR61 and 50%, *p* <0.01 for CTGF) (**Figure 7A**). In contrast, YAP1 knockdown had no effect on both downstream proteins. In GIST-882 cells, TAZ silencing alone did not significantly reduce CYR61 (*p* = ns) but a significant reduction in protein expression was observed for CTGF (45%, *p* <0.05), while YAP1 knockdown had no effect (*p* = ns versus WRAP5:siNEG). Again, dual silencing significantly reduced both gene expressions (50%, *p* <0.01 for CYR61 and 65%, *p* <0.01 for CTGF) (**Figure 7B**).

These results suggested that although TAZ appeared to play a predominant role, YAP1 could potently contribute in a complementary way in GIST-430 and GIST-882 cells. As we observed a higher basal YAP1 expression (**Figure S6**), it is probably possible that YAP1 has a more predominant role in these two GIST cell lines . Given the variability between three cell lines, both factors should be considered when targeting the YAP1/TAZ axis in GIST.

### CYR61 and CTGF as key downstream effectors mediating TAZ-induced proliferation in GIST-T1 cells

We decided to silence CYR61 and CTGF to ensure that they were effectors of the impact on cell proliferation observed for TAZ. For each gene, we designed two siRNA. Thier specific knockdown efficiency and their potent cytotoxic effect were assessed by Western blot and by LDH assay, respectively (**Figures S8A-H**). Selected siCYR61 and siCTGF led to 90% and 60% of protein inhibition at 40 nM, respectively (**Figures S8A, S8E**) without any toxic effect for the cells (**Figures S8C, S8G**. The efficacy of these selected siRNA was also validated in qPCR analyses. This experiment led to significant downregulation of *CYR61* and *CTGF* mRNA levels 24 hours after treatment with WRAP5:siCYR61 and WRAP5:siCTGF nanoparticles, respectively (-60%, *p* <0.001 for both, **Figure 8A**).

**Figure 8:**
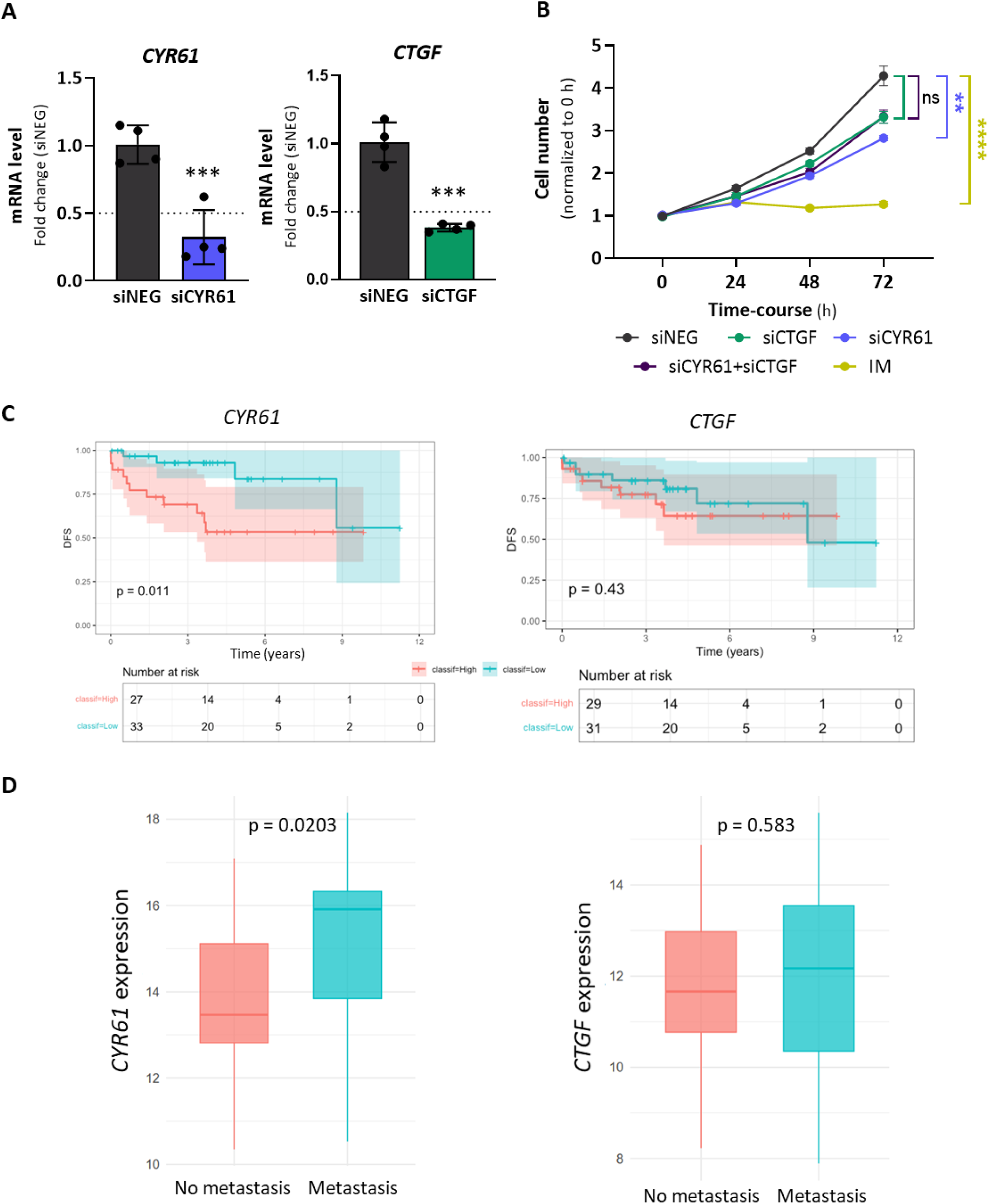
CYR61 and CTGF are effectors of TAZ for cell proliferation control. (**A**) Effect of WRAP5:siRNA nanoparticles targeting CYR61 and CTGF mRNA levels in GIST-T1 cells were analyzed by qPCR analysis 24 h post transfection. (**B**) WRAP5:siRNA nanoparticles targeting CYR61, CTGF or both (siCYR61+siCTGF) were analyzed by cell counting over a 72 h time course post-transfection (N=4 independent experiments with n=2 replicates). Abbreviation : IM = Imatinib. Conditions: WRAP5:siRNA (MR 20) with [siCYR61], [siCTGF] = 20 nM and [siNEG], [siCYR61/siCTGF] = 40 nM, siNEG was used as the control condition, IM = 50 nM. Data are presented as mean ± SD (**A**) or mean ± SEM (**B**). Statistical analyses were performed using unpaired two-tailed Student’s t-test *versus* siNEG (**A**) or using Kruskal-Wallis followed by Dunn’s multiple comparisons *versus* siNEG (**B**), ns >0.05, ** <0.01, *** <0.001, **** <0.0001. (**C**) Kaplan-Meier curves of disease-free survival (DFS) from 60 total GIST patients in the ATGsarc microarray database, stratified by *CYR61* or *CTGF* expression levels. Group 1 (red): patients with high gene expression (number initial of patients = 27) ; Group 2 (blue): patients with low gene expression (number initial of patients = 33). Number at risk = number of patients at risk over time. They are resumed in the table below the curves. (**D**) Correlation between *CYR61* or *CTGF* transcript levels and the presence of metastasis. n = 45 (red) and n = 15 (blue) in no metastatic and metastatic groups, respectively. p-values from the log-rank test are indicated.

To assess the functional impact of CYR61 and CTGF silencing on cell proliferation, we performed cell counting assays over a 72-hour time-course following WRAP5:siRNA delivery. Following GIST-T1 cellular transfection with WRAP5-based nanoparticles, we observed a downward trend in the number of cells at 48 hours or 72 hours post-transfection when CTGF was silenced alone or in combination with CYR61, respectively (around -22% at 72h, *p*= ns)(**Figure 8B**). Interstingly, CYR61 silencing alone resulted in a statistically significant reduction in cell proliferation (-34% at 72h, *p*<0.01) compared to the siNEG control.

Finally, to explore the clinical relevance of these findings, we analyzed the same cohort of 60 patients from the ATGsarc microarray database [53] as previously performed for YAP1 and TAZ (**Figure 1**). Kaplan-Meier survival analyses stratified patients by high versus low gene expression levels of *CYR61* or *CTGF*. High *CYR61* expression was significantly associated with reduced disease-free survival (*p* = 0.011; **Figure 8C**) and with the presence of metastasis during GIST evolution (*p* = 0.0203; **Figure 8D**). In contrast, *CTGF* expression showed no significant correlation with either disease-free survival (*p* = 0.43) or metastatic status (*p* = 0.583). *CYR61* is identified as a clinically relevant effector of TAZ, linking its overexpression to both increased tumor cell proliferation and poor clinical outcomes in GIST.

These results support the role of CYR61 as downstream effector of TAZ in GIST-T1 cells, to promote tumor cell proliferation. This correlates with a high *CYR61* expression associated with critical clinical features, including shorter progression-free survival and metastatic status.

## DISCUSSION

In this study, we provided the first investigation of the individual roles of YAP1 and TAZ, the two Hippo pathway effectors, in KIT-dependent IM-sensitive GIST cells. Indeed, it is well known that YAP1 and TAZ are implicated in cancer development and progression, even if most publications presented YAP1/TAZ as functionally redundant due to their structural similarity and shared transcriptional partners [24,34,35]. However, some reports suggest that both proteins have distinct roles as reviewed by Reggiani, *et al*. [37].

We aimed to evaluate whether it was possible to target YAP1 or TAZ separately using interfering RNA to elucidate which of these oncogenic proteins participated in GIST development and proliferation. Using our WRAP5-based nanoparticles as a delivery tool previously validated for therapeutic delivery [43–46], we vectorized specifically developed siRNA targeting the mRNA of *YAP1* and *TAZ*, individually, but also together using a siRNA cocktail containing an equimolar combination of both (siYAP1+siTAZ).

With this tool in our hand, we observed a specific and significant silencing of YAP1 or TAZ individually or in combination in KIT-dependent IM-sensitive GIST-T1 (**Figure 3**). However, only the silencing of TAZ resulted in a reduction in cell migration and cell proliferation (**Figure 5**). More interestingly, transcriptional activation of downstream targets such as *CYR61* and *CTGF* genes was shown after TAZ silencing, while YAP1 exerted limited or nearly no effects on these genes implicated in cell proliferation. Notably, inhibition of TAZ alone prevented the induction of oncogenic characteristics, and dual silencing showed only a slight additive effect on phenotypic outcomes. These results thus reinforce the idea of non-redundant roles in GIST cells and a more complex regulatory mechanism.

The functional divergence between YAP1 and TAZ protein function observed here was consistent with reports in other malignancies especially cancers. In hepatocellular carcinoma and cholandiocarcinoma, only YAP1 played crucial roles in the development and progression [62,63]. In non-small cell lung cancer, TAZ preferentially regulated genes involved in extracellular matrix remodeling and metastasis, whereas YAP1 controlled proliferation-related genes [64]. Moreover, the activation of TAZ nuclear translocation was associated with the highly aggressive in triple-negative subtype of breast cancer [65]. Importantly, such divergence has also been described in sarcomas, a relevant context for GIST. For example, two research articles revealed an upregulated YAP1 expression level in osteosarcoma and participated to cell differentiation [41,66], even if the role of TAZ was not evaluated. In contrast, TAZ activity was modulated for the myogenic differentiation in rhabdomyosarcoma without the participation of YAP1 [67]. Our study extended this paradigm to GIST, establishing TAZ as a main oncogenic driver and supporting the notion of tissue- and lineage-specific YAP1/TAZ activity. However, our results supported the idea that TAZ could represent a more relevant therapeutic target in the context of GIST, in line with the clinical data identifying TAZ as a superior prognostic marker (**Figure 1**).

Despite these findings, the mechanistic basis of this functional separation remained unclear as both proteins, YAP1 and TAZ, mainly bound to TEAD transcription factors to regulate gene expression involved in cancer development. This raised the possibility that differential nuclear co-factor recruitment (co-activators or co-repressors) or divergent affinity for specific TEAD isoforms could underlie the selective transcriptional activity of TAZ [37,60]. Further research should help to define these regulatory mechanisms, including chromatin context, transcriptional partner preferences, or post-translational modifications, which could fine-tune TEAD-dependent transcriptional output.

By comparing in detail the three analyzed GIST cell lines, we observed heterogeneous YAP1 and TAZ protein expression, while TAZ emerged as the dominant effector for downstream *CYR61* and *CTGF* gene downregulation (**Figures 6A, 6B, 6D, 7**). However, depending on the cellular YAP1/TAZ expression profile, a combined silencing of both proteins further repressed downstream gene expression. This suggested a possible context-dependent or co-operative role of YAP1/TAZ. This heterogeneity in YAP1/TAZ protein expression could reflect the diversity found within tumors in GIST patients and highlighted the possible advantage of targeting both pathways in more aggressive or advanced cases. Indeed, similar strategies have been effective in other cancers. In KRAS G12C mutant cells, dual knockdown of YAP1 and TAZ maximized reduction in CYR61 expression and sensitized tumors to targeted therapies [68]. Likewise, in BRAF inhibitor-resistant melanoma, dual YAP1/TAZ silencing restored drug sensitivity and suppressed proliferation [69,70]. These findings supported our conclusion that dual inhibition, while not always synergistic, could provide broader efficacy in complex tumor environments.

In the present study, it was demonstrated that upon Hippo inactivation, the TAZ-CYR61 axis is activated as evidenced by the proliferation assay (**Figure 8B**). This activation exerts a negative impact on clinical progression as shown by correlating TAZ/CYR61 expression in the GIST database for progression-free survival and metastasis formation (**Figures 8C, 8D**). Indeed, the two genes are well established downstream targets of the YAP1/TAZ-TEAD complex and are implicated in cell proliferation, angiogenesis, extracellular matrix deposition, and metastasis across multiple tumor types [24,35].

In the three used GIST cell models, their expression was consistently regulated by TAZ and less by YAP1, reinforcing the specificity of TAZ-mediated transcription in GIST. Similar finding was revealed in gastric cancer, another digestive cancer, showing that TAZ upregulation was an important inducer of epithelial-mesenchymal transition and gastric cancer stem cell [55].

Although CD1 expression remained unaffected when YAP1 and TAZ were inhibited, this does not exclude the involvement of other cell-cycle regulators such as cyclin D2, cyclin D3, CDK2, or CDK4 notably in TAZ-driven proliferation [71]. Moreover, a modulation of apoptosis-related markers such as MCL-1 or BCL-2 could not be disregarded, leading to an indirect impact through other survival pathways [72]. Our results suggest that YAP1 and TAZ silencing does not significantly alter the major downstream signaling pathways of KIT activation, ERK/MAPK and PI3K/AKT/mTOR, indicating that TAZ-driven transcriptional programs may promote tumor progression through pathways at least partly independent of canonical KIT signaling.

In conclusion, the use of WRAP5:siRNA nanoparticles could be an innovative therapeutic strategy for GIST treatment, as these nanoparticles exhibited low cytotoxicity and enabled sustained delivery of siRNA *in vitro* and *in vivo* as previously reported in other pathologies or in GIST [42,44,46–48,73]. This approach opened new avenues for RNA-based therapeutics in GIST, where current treatment options remained limited for GIST patients to kinase inhibition using TKI [74]. Notably, siRNA-based drugs are now commonly accepted as six vectorized siRNA have been approved by the FDA (US Food and Drug Administration) and the EMA (European Medicines Agency) [75]. In the context of the presented siTAZ or (siYAP1+siTAZ)-loaded WRAP5-based nanoparticles as therapeutic entity, further *in vivo* investigations will be needed to further promote their development. Especially, further work should be performed in order to determine if both siRNA should be used to overcome GIST patient heterogenicity (variation in the mutation profile).

Finally, to potentiate siRNA-loaded WRAP5 nanoparticles, other parameters should be also taken in account such as the optimization of the nanoparticle itself through PEGylation (increasing the stability) or targeting motif insertion as reported previously in other physio-pathological contexts [46,48,73]. Given emerging links between Hippo pathway reactivation and TKI resistance [18], future work should also explore how YAP1/TAZ modulation impacts IM response and resistance mechanisms.

## SUPPPLEMENTARY INFORMATION

Supplementary Material and Figures

## ETHICS APPROVAL AND CONSENT TO PARTICIPATE

Not applicable.

## CONSENT FOR PUBLICATION

Not applicable

## AVAILABILITY OF DATA AND MATERIALS

GIST data base (ATGsarc) is available at https://atg-sarc.sarcomabcb.org/

## AUTHORS’ CONTRIBUTIONS

Conceptualization, I.P., P.B., S.D., P.D.S.B., S.F., C.S.; Methodology, I.P., P.B., S.D., E.V., F.C; Experiments, I.P., P.B., S.D.; Figures, I.P., P.B., S.D., F.C; Statistical analysis, I.P., P.B., F.C; Writing-original draft preparation, I.P., P.B., S.D., S.F., P.D.S.B., C.S., E.V., F.C.; Funding acquisition, P.B., S.F., P.D.S.B. All authors have read and agreed to the published version of the manuscript.

## ACKNOWLEDGMENTS

The authors are grateful to Pascal Verdié from the SynBio3 platform for providing peptide synthesis facilities, and to Pierre Sanchez for performing LCMS analysis, both from the Institut des Biomolécules Max Mousseron (IBMM), Montpellier (France). The authors thank Dr. Karidia Konate and Thanian Hammoum for critical reading of the manuscript.

## REFERENCES

1. Blay J-Y, Kang Y-K, Nishida T, von Mehren M. Gastrointestinal stromal tumours. Nat Rev Dis Primer. 2021;7:22.

2. Casali PG, Blay JY, Abecassis N, Bajpai J, Bauer S, Biagini R, et al. Gastrointestinal stromal tumours: ESMO-EURACAN-GENTURIS Clinical Practice Guidelines for diagnosis, treatment and follow-up. Ann Oncol Off J Eur Soc Med Oncol. 2022;33:20–33.

3. Min KW, Leabu M. Interstitial cells of Cajal (ICC) and gastrointestinal stromal tumor (GIST): facts, speculations, and myths. J Cell Mol Med. 2006;10:995–1013.

4. Hirota S, Isozaki K, Moriyama Y, Hashimoto K, Nishida T, Ishiguro S, et al. Gain-of-function mutations of c-kit in human gastrointestinal stromal tumors. Science. 1998;279:577–80.

5. Schaefer I-M, DeMatteo RP, Serrano C. The GIST of Advances in Treatment of Advanced Gastrointestinal Stromal Tumor. Am Soc Clin Oncol Educ Book. 2022;42:1–15.

6. Bauer S, George S, von Mehren M, Heinrich MC. Early and Next-Generation KIT/PDGFRA Kinase Inhibitors and the Future of Treatment for Advanced Gastrointestinal Stromal Tumor. Front Oncol. 2021;11.

7. Serrano C, George S. Gastrointestinal Stromal Tumor: Challenges and Opportunities for a New Decade. Clin Cancer Res Off J Am Assoc Cancer Res. 2020;26:5078–85.

8. Duan Y, Haybaeck J, Yang Z. Therapeutic Potential of PI3K/AKT/mTOR Pathway in Gastrointestinal Stromal Tumors: Rationale and Progress. Cancers. 2020;12:2972.

9. Zhou S, Abdihamid O, Tan F, Zhou H, Liu H, Li Z, et al. KIT mutations and expression: current knowledge and new insights for overcoming IM resistance in GIST. Cell Commun Signal. 2024;22:153.

10. Demetri GD, von Mehren M, Blanke CD, Van den Abbeele AD, Eisenberg B, Roberts PJ, et al. Efficacy and safety of imatinib mesylate in advanced gastrointestinal stromal tumors. N Engl J Med. 2002;347:472–80.

11. Demetri GD, Oosterom AT van, Garrett CR, Blackstein ME, Shah MH, Verweij J, et al. Efficacy and safety of sunitinib in patients with advanced gastrointestinal stromal tumour after failure of imatinib: a randomised controlled trial. The Lancet. 2006;368:1329–38.

12. Demetri GD, Reichardt P, Kang Y-K, Blay J-Y, Rutkowski P, Gelderblom H, et al. Efficacy and safety of regorafenib for advanced gastrointestinal stromal tumours after failure of imatinib and sunitinib (GRID): an international, multicentre, randomised, placebo-controlled, phase 3 trial. The Lancet. 2013;381:295–302.

13. Serrano C, Fletcher JA. Overcoming heterogenity in imatinib-resistant gastrointestinal stromal tumor. Oncotarget. 2019;10:6286–7.

14. Chen T, Ni N, Yuan L, Xu L, Bahri N, Sun B, et al. Proteasome Inhibition Suppresses KIT-Independent Gastrointestinal Stromal Tumors Via Targeting Hippo/YAP/Cyclin D1 Signaling. Front Pharmacol. 2021;12:686874.

15. Ou W-B, Ni N, Zuo R, Zhuang W, Zhu M, Kyriazoglou A, et al. Cyclin D1 is a mediator of gastrointestinal stromal tumor KIT-independence. Oncogene. 2019;38:6615–29.

16. Tu Y, Zuo R, Ni N, Eilers G, Wu D, Pei Y, et al. Activated tyrosine kinases in gastrointestinal stromal tumor with loss of KIT oncoprotein expression. Cell Cycle Georget Tex. 2018;17:2577–92.

17. Guérin A, Martire D, Trenquier E, Lesluyes T, Sagnol S, Pratlong M, et al. LIX1 regulates YAP activity and controls gastrointestinal cancer cell plasticity. J Cell Mol Med. 2020;24:9244–54.

18. Ruiz-Demoulin S, Trenquier E, Dekkar S, Deshayes S, Boisguérin P, Serrano C, et al. LIX1 Controls MAPK Signaling Reactivation and Contributes to GIST-T1 Cell Resistance to Imatinib. Int J Mol Sci. 2023;24:7138.

19. Guérin A, Angebault C, Kinet S, Cazevieille C, Rojo M, Fauconnier J, et al. LIX1-mediated changes in mitochondrial metabolism control the fate of digestive mesenchyme-derived cells. Redox Biol. 2022;56:102431.

20. Fu M, Hu Y, Lan T, Guan K-L, Luo T, Luo M. The Hippo signalling pathway and its implications in human health and diseases. Signal Transduct Target Ther. 2022;7:376.

21. Ma S, Meng Z, Chen R, Guan K-L. The Hippo Pathway: Biology and Pathophysiology. Annu Rev Biochem. 2019;88:577–604.

22. Boopathy GTK, Hong W. Role of Hippo Pathway-YAP/TAZ Signaling in Angiogenesis. Front Cell Dev Biol. 2019;7.

23. LeBlanc L, Ramirez N, Kim J. Context-dependent roles of YAP/TAZ in stem cell fates and cancer. Cell Mol Life Sci CMLS. 2021;78:4201–19.

24. Messina B, Lo Sardo F, Scalera S, Memeo L, Colarossi C, Mare M, et al. Hippo pathway dysregulation in gastric cancer: from Helicobacter pylori infection to tumor promotion and progression. Cell Death Dis. 2023;14:1–12.

25. Mouillet-Richard S, Laurent-Puig P. YAP/TAZ Signalling in Colorectal Cancer: Lessons from Consensus Molecular Subtypes. Cancers. 2020;12:3160.

26. Montalto FI, De Amicis F. Cyclin D1 in Cancer: A Molecular Connection for Cell Cycle Control, Adhesion and Invasion in Tumor and Stroma. Cells. 2020;9:2648.

27. Qie S, Diehl JA. Cyclin D1, Cancer Progression and Opportunities in Cancer Treatment. J Mol Med Berl Ger. 2016;94:1313–26.

28. Wu X, Yamashita K, Lou M, Matsumoto C, Zhang W, Baba H, et al. AT101 Suppresses Gastrointestinal Stromal Tumor Growth and Promotes Apoptosis via YAP/TAZ-CCND1 and FBXW7-MCL1 Axes. Ann Surg Oncol. 2025;32:5991–6004.

29. Delvaux M, Hagué P, Craciun L, Wozniak A, Demetter P, Schöffski P, et al. Ferroptosis Induction and YAP Inhibition as New Therapeutic Targets in Gastrointestinal Stromal Tumors (GISTs). Cancers. 2022;14:5050.

30. Vandenberghe P, Delvaux M, Hagué P, Erneux C, Vanderwinden J-M. Potentiation of imatinib by cilostazol in sensitive and resistant gastrointestinal stromal tumor cell lines involves YAP inhibition. Oncotarget. 2019;10:1798–811.

31. Moroishi T, Hansen CG, Guan K-L. The emerging roles of YAP and TAZ in cancer. Nat Rev Cancer. 2015;15:73–9.

32. Driskill JH, Dermawan JK, Antonescu CR, Pan D. YAP, TAZ, and Hippo-Dysregulating Fusion Proteins in Cancer. Annu Rev Cancer Biol. 2024;8:331–50.

33. Wei Y, Hui VLZ, Chen Y, Han R, Han X, Guo Y. YAP/TAZ: Molecular pathway and disease therapy. MedComm. 2023;4:e340.

34. Cunningham R, Hansen CG. The Hippo pathway in cancer: YAP/TAZ and TEAD as therapeutic targets in cancer. Clin Sci Lond Engl 1979. 2022;136:197–222.

35. Luo J, Deng L, Zou H, Guo Y, Tong T, Huang M, et al. New insights into the ambivalent role of YAP/TAZ in human cancers. J Exp Clin Cancer Res CR. 2023;42:130.

36. Piccolo S, Panciera T, Contessotto P, Cordenonsi M. YAP/TAZ as master regulators in cancer: modulation, function and therapeutic approaches. Nat Cancer. 2023;4:9–26.

37. Reggiani F, Gobbi G, Ciarrocchi A, Sancisi V. YAP and TAZ Are Not Identical Twins. Trends Biochem Sci. 2021;46:154–68.

38. Bae JS, Kim SM, Lee H. The Hippo signaling pathway provides novel anti-cancer drug targets. Oncotarget. 2016;8:16084–98.

39. Chen C, Zhou H, Zhang X, Liu Z, Ma X. PRMT1 potentiates chondrosarcoma development through activation of YAP activity. Mol Carcinog. 2019;58:2193–206.

40. Patterson MR, Cogan JA, Cassidy R, Theobald DA, Wang M, Scarth JA, et al. The Hippo pathway transcription factors YAP and TAZ play HPV-type dependent roles in cervical cancer. Nat Commun. 2024;15:5809.

41. Zhang Y-H, Li B, Shen L, Shen Y, Chen X-D. The role and clinical significance of YES-associated protein 1 in human osteosarcoma. Int J Immunopathol Pharmacol. 2013;26:157–67.

42. Konate K, Dussot M, Aldrian G, Vaissière A, Viguier V, Neira IF, et al. Peptide-Based Nanoparticles to Rapidly and Efficiently “Wrap ‘n Roll” siRNA into Cells. Bioconjug Chem. 2019;30:592–603.

43. Konate K, Josse E, Tasic M, Redjatti K, Aldrian G, Deshayes S, et al. WRAP-based nanoparticles for siRNA delivery: a SAR study and a comparison with lipid-based transfection reagents. J Nanobiotechnology. 2021;19:236.

44. Konate K, Pezzati I, Redjatti K, Agnel E, Vivès E, Faure S, et al. Multiprotein Silencing Using WRAP-Based Nanoparticles: A Proof of Concept. Bioconjug Chem. 2025;36:1218–33.

45. Boisguérin P, Konate K, Josse E, Vivès E, Deshayes S. Peptide-Based Nanoparticles for Therapeutic Nucleic Acid Delivery. Biomedicines. 2021;9:583.

46. Di Gregorio G, Vallée C, Konate K, Teko-Agbo CA, Hammoum T, Faure-Gautron H, et al. Enhancing WRAP-Based Nanoparticles for Small Interfering Ribonucleic Acid Delivery in pH-Sensitive Environments. ChemMedChem. 2025;e2400885.

47. Ferreiro I, Genevois C, Konate K, Vivès E, Boisguérin P, Deshayes S, et al. In Vivo Follow-Up of Gene Inhibition in Solid Tumors Using Peptide-Based Nanoparticles for siRNA Delivery. Pharmaceutics. 2021;13:749.

48. Konate K, Teko-Agbo CA, Pezzati I, Hammoum T, Deshayes S, Descamps S, et al. WRAP-based nanoparticles for siRNA delivery in zebrafish embryos by simple bath immersion. Mol Ther Methods Clin Dev. 2025;33:101458.

49. García-Valverde A, Rosell J, Serna G, Valverde C, Carles J, Nuciforo P, et al. Preclinical Activity of PI3K Inhibitor Copanlisib in Gastrointestinal Stromal Tumor. Mol Cancer Ther. 2020;19:1289–97.

50. Taguchi T, Sonobe H, Toyonaga S, Yamasaki I, Shuin T, Takano A, et al. Conventional and molecular cytogenetic characterization of a new human cell line, GIST-T1, established from gastrointestinal stromal tumor. Lab Investig J Tech Methods Pathol. 2002;82:663–5.

51. Lux ML, Rubin BP, Biase TL, Chen C-J, Maclure T, Demetri G, et al. KIT Extracellular and Kinase Domain Mutations in Gastrointestinal Stromal Tumors. Am J Pathol. 2000;156:791–5.

52. Garner AP, Gozgit JM, Anjum R, Vodala S, Schrock A, Zhou T, et al. Ponatinib inhibits polyclonal drug-resistant KIT oncoproteins and shows therapeutic potential in heavily pretreated gastrointestinal stromal tumor (GIST) patients. Clin Cancer Res Off J Am Assoc Cancer Res. 2014;20:5745–55.

53. Lagarde P, Pérot G, Kauffmann A, Brulard C, Dapremont V, Hostein I, et al. Mitotic checkpoints and chromosome instability are strong predictors of clinical outcome in gastrointestinal stromal tumors. Clin Cancer Res Off J Am Assoc Cancer Res. 2012;18:826–38.

54. McKey J, Martire D, de Santa Barbara P, Faure S. LIX1 regulates YAP1 activity and controls the proliferation and differentiation of stomach mesenchymal progenitors. BMC Biol. 2016;14:34.

55. Tiffon C, Giraud J, Molina-Castro SE, Peru S, Seeneevassen L, Sifré E, et al. TAZ Controls Helicobacter pylori-Induced Epithelial-Mesenchymal Transition and Cancer Stem Cell-Like Invasive and Tumorigenic Properties. Cells. 2020;9:1462.

56. Burkhardt F, Spies BC, Wesemann C, Schirmeister CG, Licht EH, Beuer F, et al. Cytotoxicity of polymers intended for the extrusion-based additive manufacturing of surgical guides. Sci Rep. 2022;12:7391.

57. Pavel M, Renna M, Park SJ, Menzies FM, Ricketts T, Füllgrabe J, et al. Contact inhibition controls cell survival and proliferation via YAP/TAZ-autophagy axis. Nat Commun. 2018;9:2961.

58. Zhang Q, Han X, Chen J, Xie X, Xu J, Zhao Y, et al. Yes-associated protein (YAP) and transcriptional coactivator with PDZ-binding motif (TAZ) mediate cell density–dependent proinflammatory responses. J Biol Chem. 2018;293:18071–85.

59. Ghaboura N. Unraveling the Hippo pathway: YAP/TAZ as central players in cancer metastasis and drug resistance. EXCLI J. 2025;24:612–37.

60. Zanconato F, Cordenonsi M, Piccolo S. YAP/TAZ at the Roots of Cancer. Cancer Cell. 2016;29:783–803.

61. Mao X, Yang X, Chen X, Yu S, Yu S, Zhang B, et al. Single-cell transcriptome analysis revealed the heterogeneity and microenvironment of gastrointestinal stromal tumors. Cancer Sci. 2021;112:1262–74.

62. Li S, Hao L, Li N, Hu X, Yan H, Dai E, et al. Targeting the Hippo/YAP1 signaling pathway in hepatocellular carcinoma: From mechanisms to therapeutic drugs (Review). Int J Oncol. 2024;65:1–17.

63. Pei T, Li Y, Wang J, Wang H, Liang Y, Shi H, et al. YAP is a critical oncogene in human cholangiocarcinoma. Oncotarget. 2015;6:17206–20.

64. Shreberk-Shaked M, Dassa B, Sinha S, Di Agostino S, Azuri I, Mukherjee S, et al. A Division of Labor between YAP and TAZ in Non-Small Cell Lung Cancer. Cancer Res. 2020;80:4145–57.

65. Díaz-Martín J, López-García MÁ, Romero-Pérez L, Atienza-Amores MR, Pecero ML, Castilla MÁ, et al. Nuclear TAZ expression associates with the triple-negative phenotype in breast cancer. Endocr Relat Cancer. 2015;22:443–54.

66. Basu-Roy U, Han E, Rattanakorn K, Gadi A, Verma N, Maurizi G, et al. PPARγ agonists promote differentiation of cancer stem cells by restraining YAP transcriptional activity. Oncotarget. 2016;7:60954–70.

67. Hebron KE, Perkins OL, Kim A, Jian X, Girald-Berlingeri SA, Lei H, et al. ASAP1 and ARF1 Regulate Myogenic Differentiation in Rhabdomyosarcoma by Modulating TAZ Activity. Mol Cancer Res. 2025;23:95–106.

68. Edwards AC, Stalnecker CA, Jean Morales A, Taylor KE, Klomp JE, Klomp JA, et al. TEAD Inhibition Overcomes YAP1/TAZ-Driven Primary and Acquired Resistance to KRASG12C Inhibitors. Cancer Res. 2023;83:4112–29.

69. Kim H, Son S, Ko Y, Lee JE, Kim S, Shin I. YAP, CTGF and Cyr61 are overexpressed in tamoxifen-resistant breast cancer and induce transcriptional repression of ERα. J Cell Sci. 2021;134:jcs256503.

70. Kim MH, Kim J, Hong H, Lee S, Lee J, Jung E, et al. Actin remodeling confers BRAF inhibitor resistance to melanoma cells through YAP/TAZ activation. EMBO J. 2016;35:462–78.

71. Knapp CM, He J, Lister J, Whitehead KA. Lipid nanoparticle siRNA cocktails for the treatment of mantle cell lymphoma. Bioeng Transl Med. 2018;3:138–47.

72. Wang L, Chen Y, Zhang M, Liu J, Li H, Liu M, et al. Chemical dissection of selective myeloid leukemia-1 inhibitors: How they were found and evolved. Eur J Med Chem. 2025;283:117168.

73. Faria R, Vivès E, Boisguérin P, Descamps S, Sousa Â, Costa D. Upgrading Mitochondria-Targeting Peptide-Based Nanocomplexes for Zebrafish In Vivo Compatibility Assays. Pharmaceutics. 2024;16:961.

74. Joensuu H, Miyashita H, George S, Sicklick J. Navigating Ongoing Challenges in GI Stromal Tumors. Am Soc Clin Oncol Educ Book Am Soc Clin Oncol Annu Meet. 2025;45:e473224.

75. Tang Q, Khvorova A. RNAi-based drug design: considerations and future directions. Nat Rev Drug Discov. 2024;23:341–64.

