## Supplemental data for "Breaking the redundancy: TAZ outperforms YAP1 in GIST progression"

**SUPPORTING INFORMATION**

**Supporting Tables**

**Table S1: Used siRNA sequences**

| ID | Sense sequence | Anti-sense sequence | Reference |
|---|---|---|---|
| **siYAP1** | 5’-CUA-UGU-UCA-UUC-CAU-CUC-CdTdT-3’ | 5’-GGA-GAU-GGA-AUG-AAC-AUA-GdTdT-3’ | This work |
| **siYAP1’** | 5’-AUA-GUA-AAU-UUC-UCC-AUC-CdTdT-3’ | 5’-GGA-UGG-AGA-AAU-UUA-CUA-UdTdT-3’ | This work |
| **siTAZ** | 5’-AAA-UCA-GGG-AAA-CGG-GUC-UdTdT-3’ | 5’-AGA-CCC-GUU-UCC-CUG-AUU-UdTdT-3’ | This work |
| **siTAZ’** | 5’-AUU-AUU-AGU-GAU-GGA-UCU-CdTdT-3’ | 5’-GAG-AUC-CAU-CAC-UAA-UAA-UdTdT-3’ | This work |
| **si(YAPTAZ)** | 5’-UGU-GGA-UGA-GAU-GGA-UAC-AdTdT-3’ | 5’-UGU-AUC-CAU-CUC-AUC-CAC-AdTdT-3’ | (Tiffon et al., 2020) |
| **siCYR61** | 5’-AUC-AUC-AUG-ACG-UUC-UUG-GdTdT-3’ | 5’-CCA-AGA-ACG-UCA-UGA-UGA-UdTdT-3’ | This work |
| **siCYR61’** | 5’-UGG-AGU-UGA-CGA-GAA-ACA-AdTdT-3’ | 5’-UUG-UUU-CUC-GUC-AAC-UCC-AdTdT-3’ | This work |
| **siCTGF** | 5'-AAU-UUA-GCU-CGG-UAU-GUC-UdTdT-3' | 5'-AGA-CAU-ACC-GAG-CUA-AAU-UdTdT-3' | This work |
| **siCTGF’** | 5’-AUU-UUG-GGA-GUA-CGG-AUG-CpTpT-3’ | 5’-GCA-UCC-GUA-CUC-CCA-AAA-UpTpT-3’ | |
| **siNEG** | Confidential | Confidential | Negative Control siRNA, Eurogentec, Réf. SR-CL000-005 |

**Table S2: List of antibodies**

| Target (Protein) | MW (kDa) | Host Species | Clone / Catalog Number | Dilution | Company / Supplier | Application |
|---|---|---|---|---|---|---|
| **YAP1** | 65 - 78 | Rabbit | D8H1X / #14074 | 1:1,000 (GIST-T1) 1:5,000 (GIST-882) 1:10,000 (GIST-430) | Cell Signaling | WB |
| **YAP1** | 65-70 | Mouse | 66900-1 | 1:800 | Proteintech | If |
| **TAZ** | 55 | Rabbit | E9J5A / #72804 | 1:1,000 (GIST-T1 and GIST-882) 1:5,000 (GIST-430) 1:500 (If) | Cell Signaling | WB, If |
| **KIT (CD117)** | 145 | Rabbit | #DB062 | 1:1,000 | DB Biotech | WB |
| **Phospho-KIT (pKIT)** | 145 | Rabbit | Tyr703 D12E12 / #3073 | 1:1,000 | Cell Signaling | WB |
| **AKT** | 60 | Rabbit | #9272 | 1:1,000 | Cell Signaling | WB |
| **Phospho-AKT (pAKT)** | 60 | Rabbit | Ser473 D9E / #4060 | 1:1,000 | Cell Signaling | WB |
| **p44/42 MAPK (Erk1/2)** | 42, 44 | Mouse | L34F12 / #4696 | 1:1,000 | Cell Signaling | WB |
| **Phospho-p44/42 MAPK (Erk1/2)** | 42, 44 | Rabbit | Thr202/Tyr204 20G11 / #4376 | 1:1,000 | Cell Signaling | WB |
| **CYR61** | 41 | Rabbit | D4H5D / #14479 | 1:1,000 | Cell Signaling | WB |
| **CTGF** | 35 | Rabbit | D8Z8U / #86641 | 1:1,000 | Cell Signaling | WB |
| **CD1** | 34 | Rabbit | SP4 / #MA5-16356 | 1:1,000 | ThermoFisher Scientific | WB |
| **VINCULIN** | 124 | Rabbit | E1E9V / #13901 | 1:1,000 | Cell Signaling | WB |
| **α-ACTININ** | 100 | Rabbit | D6F6 / #6487 | 1:1,000 | Cell Signaling | WB |
| **β-ACTIN** | 42 | Mouse | #A5441 | 1:10,000 | Sigma-Aldrich | WB |
| **Anti-Rabbit** | / | IgG, HRP | #7074 | 1:1,000 | Cell Signaling | WB |
| **Anti-Mouse** | / | IgG, HRP | #7076 | 1:10,000 (β-Actin) 1:1,000 (others) | Cell Signaling | WB |
| **Anti-Rabbit Alexa Fluor™ 488** | / | Donkey | #A31570 | 1:1,000 | Invitrogen | If |
| **Anti-Mouse Alexa Fluor™ 555** | / | Donkey | #A21206 | 1:1,000 | Invitrogen | If |

Footnotes: WB: Western Blot / If: Immunofluorescence.

**Table S3: Human gene-specific primers used for RT-qPCR.**

| Target | Forward primer (5'-3') | Reverse primer (5'-3') | Amplicon (bp) | Reference |
|---|---|---|---|---|
| **YAP1** | CACAGCATGTTCGAGCTCAT | GATGCTGAGCTGTGGGTGTA | 120 | (Molina-Castro et al., 2020) |
| **TAZ** | TGCCGTCAGTTCCACAC | GTTCTGCTGGCTCAGGGT | 115 | This work |
| **CYR61** | GGAGCCTCGCATCCTAT | ATTGGTAACTCGTGTGGAGA | 115 | (Guérin et al., 2020) |
| **CTGF** | TTCAAGTGCCCTGACGG | GCGATTCAAAGATGTCATTGTCTC | 117 | This work |
| **HPRT1** | TGGTCAGGCAGTATAATCCA | GGTCCTTTTCACCAGCAAGCT | 59 | (Molina-Castro et al., 2020) |

**REFERENCES**


1. Guérin, A., Martire, D., Trenquier, E., Lesluyes, T., Sagnol, S., Pratlong, M., Lefebvre, E., Chibon, F., de Santa Barbara, P., & Faure, S. (2020). LIX1 regulates YAP activity and controls gastrointestinal cancer cell plasticity. Journal of Cellular and Molecular Medicine, 24(16), 9244-9254. https://doi.org/10.1111/jcmm.15569
2. Molina-Castro, S. E., Tiffon, C., Giraud, J., Boeuf, H., Sifre, E., Giese, A., Belleannée, G., Lehours, P., Bessède, E., Mégraud, F., Dubus, P., Staedel, C., & Varon, C. (2020). The Hippo Kinase LATS2 Controls Helicobacter pylori-Induced Epithelial-Mesenchymal Transition and Intestinal Metaplasia in Gastric Mucosa. Cellular and Molecular Gastroenterology and Hepatology, 9(2), 257-276. https://doi.org/10.1016/j.jcmgh.2019.10.007
3. Tiffon, C., Giraud, J., Molina-Castro, S. E., Peru, S., Seeneevassen, L., Sifré, E., Staedel, C., Bessède, E., Dubus, P., Mégraud, F., Lehours, P., Martin, O. C. B., & Varon, C. (2020). TAZ Controls Helicobacter pylori-Induced Epithelial-Mesenchymal Transition and Cancer Stem Cell-Like Invasive and Tumorigenic Properties. Cells, 9(6), 1462. https://doi.org/10.3390/cells9061462

### Supporting Figures

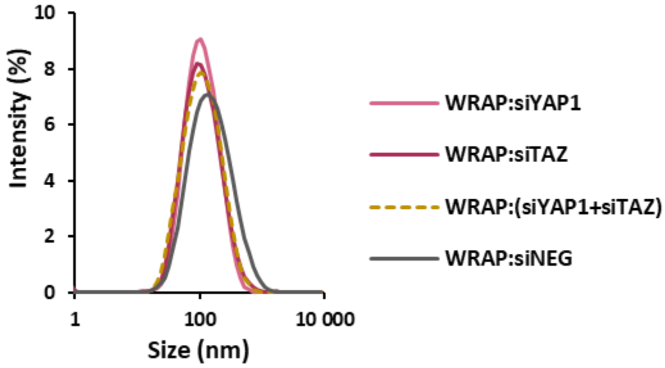


| Size (nm) | PdI |
| --- | --- |
| 92.7 ± 2.1 | 0.29 ± 0.06 |
| 92.8 ± 11.9 | 0.30 ± 0.04 |
| 89.3 ± 8.9 | 0.29 ± 0.02 |
| 121.9 ± 7.3 | 0.36 ± 0.01 |

**Figure S1: Characterization of WRAP5:siRNA nanoparticles by dynamic light scattering (DLS).**
Mean size distribution of WRAP5:siRNA nanoparticles (W5 = 10 µM, MR = 20)
DLS measurements were performed on all nanoparticles, with three individual experiments (N = 3) and three runs per condition.

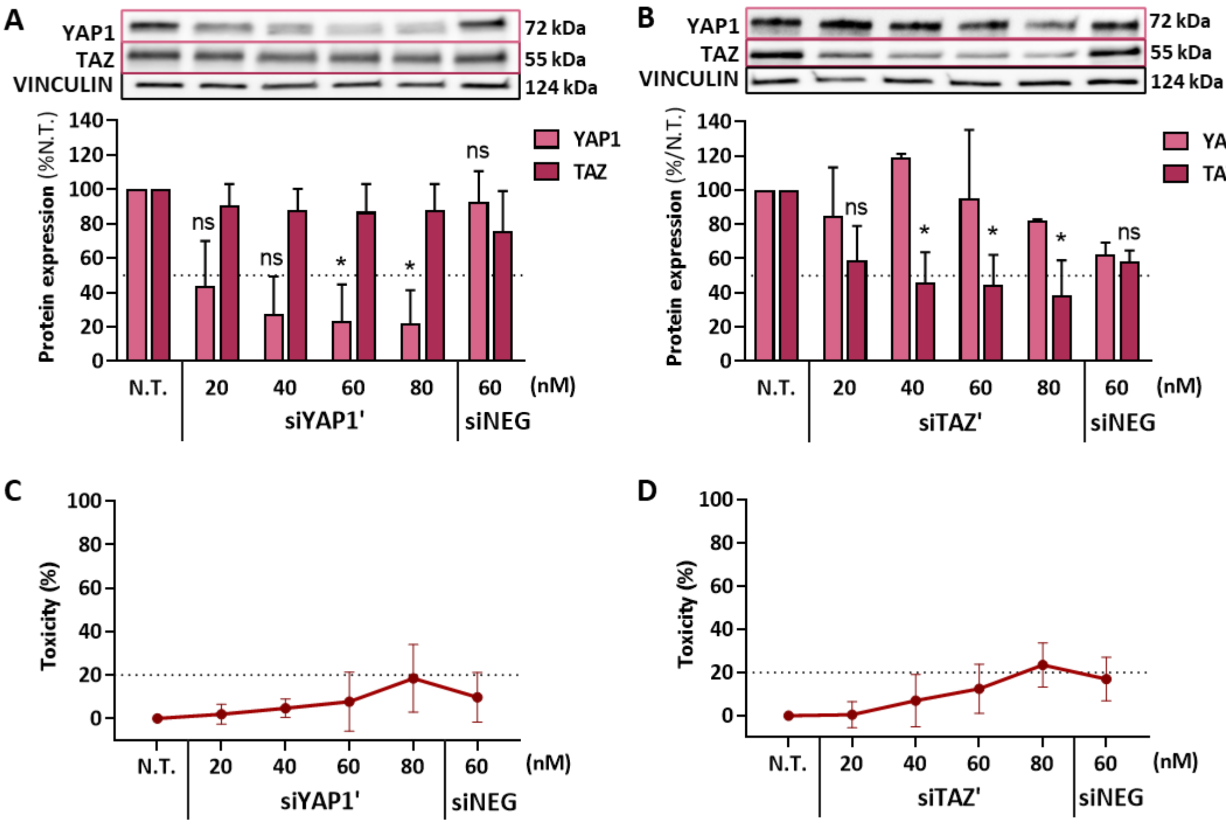


**Figure S2: Evaluation of YAP1 or TAZ silencing in GIST-T1 cells with WRAP5:siRNA' nanoparticles**
WRAP5:siRNA nanoparticles delivering siYAP1' (**A**) or siTAZ' (**B**) induced a dose-dependent inhibition of YAP1 or TAZ in GIST-T1 cells after 48 h of incubation, as shown by Western blot quantification. A slight toxicity was observed respectively for siYAP1' (**C**) and siTAZ' (**D**) at 80 nM and above, as assessed by LDH assay.
Controls include untreated (N.T.) and siNEG-treated cells. Data are presented as mean ± SD from N=4 independent experiments. Statistical analysis was performed using Kruskal-Wallis followed by Dunn's multiple comparisons test *versus* N.T. (**A**, **B**); ns >0.05, * <0.05. Analysis of TAZ expression following siYAP1 treatment and YAP1 expression following siTAZ treatment showed no significant differences compared to N.T. (data not annotated on the graph for clarity).

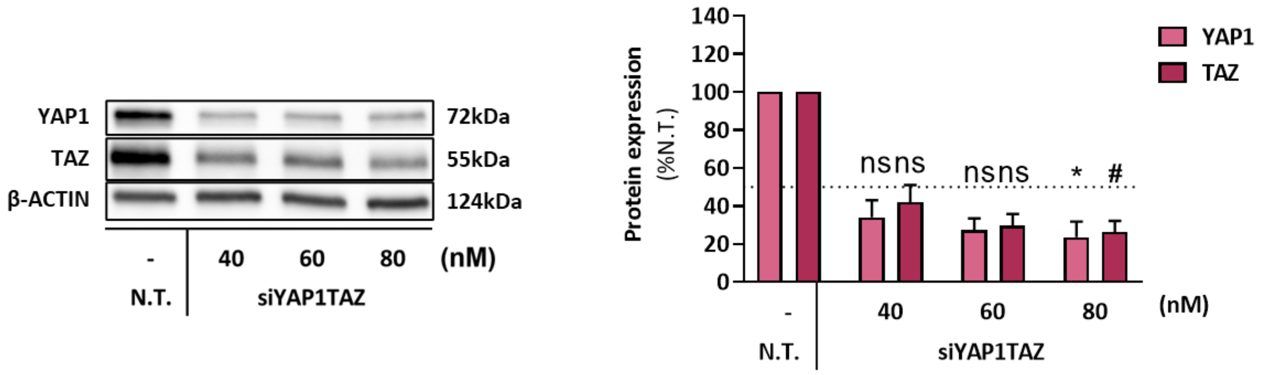


**Figure S3: Silencing YAP1 and TAZ protein expression using a single siRNA in GIST-T1 cells**
WRAP5-based nanoparticles encapsulating a single siRNA (siYAP1TAZ) published by Tiffon et al., 2020 induced a dose-dependent inhibition of YAP1 and TAZ simultaneously in GIST-T1 cells after 48 h of incubation, as shown by Western blot quantification.
Values are the mean ± SD for N=3 independent experiments. Statistical test: Kruskal-Wallis followed by Dunn with ns >0.05 and * or # <0.05 comparisons test *versus* siNEG.

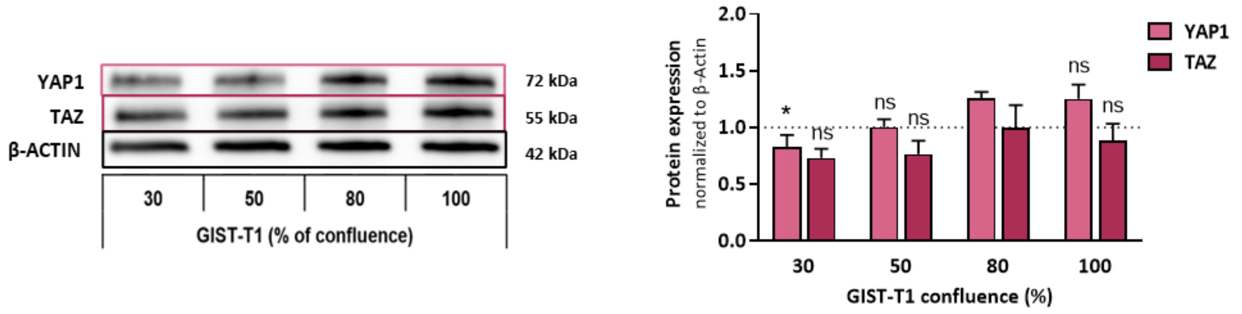


**Figure S4 : YAP1 and TAZ protein expression in GIST-T1 cells**
Western blot analysis and quantification of YAP1 and TAZ protein expression in GIST-T1 cells according to cell confluence. Values are the mean ± SEM for N=4 independent experiments. Statistical test: Kruskal-Wallis followed by Dunn with ns >0.05 and * <0.05 for YAP1 or TAZ expression at different confluences *versus* 80% confluence.

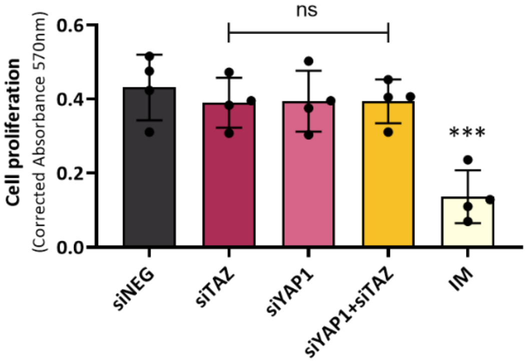


**Figure S5: Cell proliferation remains unaffected at 48 h post siRNA transfection**
Proliferation of GIST-T1 cells was assessed 48 h after transfection using the CellTiter 96® Non-Radioactive Cell Proliferation Assay for N=4 independent experiments with n=5 replicates each.
siNEG was used as a control. Abbreviation: IM = Imatinib. Data are presented as mean ± SD. Statistical analysis was performed using one-way ANOVA followed by Dunnett's multiple comparisons test *versus* siNEG; ns > 0.05, *** < 0.001.

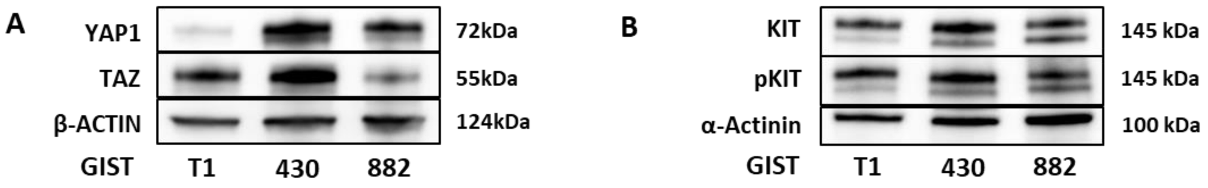


**Figure S6: Basal expression of YAP1, TAZ, KIT, pKIT in GIST cell lines.**
Comparison of the basal protein expression levels of YAP1 and TAZ (**A**) as well as KIT and phosphorylated KIT (pKIT) (**B**) in GIST-T1, GIST-430, and GIST-882 cell lines, as assessed by Western blot.

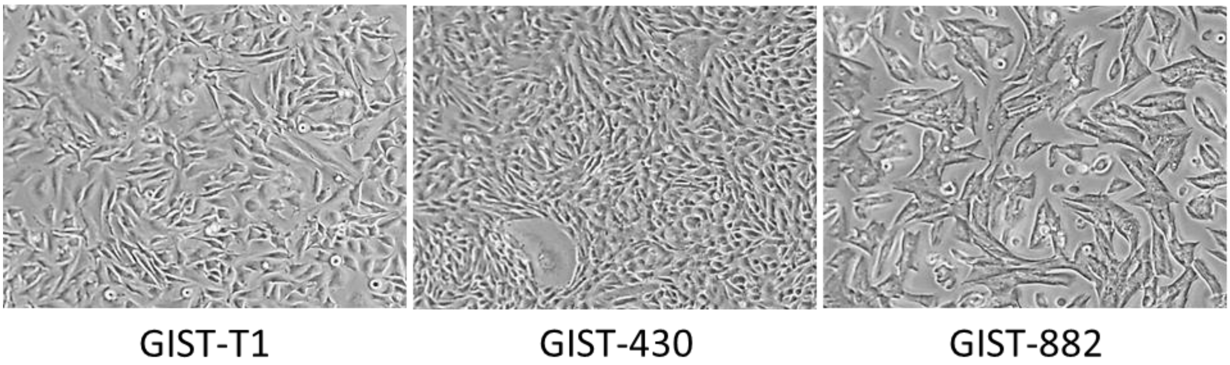


**Figure S7: Morphology of three different GIST cell lines**
GIST-T1, GIST-430, GIST-882 images taken using the EVOS® LX Core microscope system with an 10x objective (#AMEX1000, Thermo Fisher Scientific).

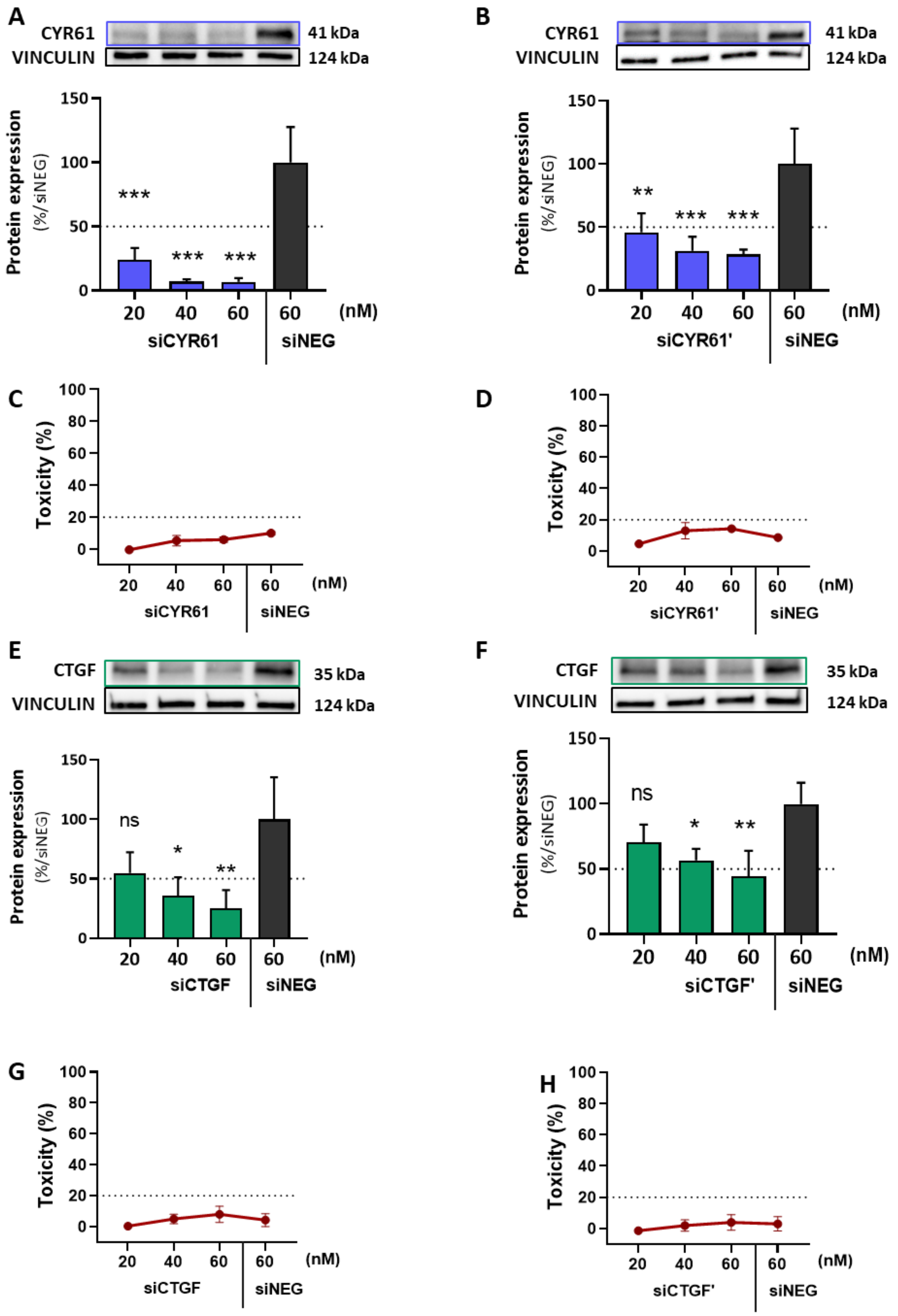


**Figure S8: Evaluation of CYR61 or CTGF silencing in GIST-T1 cells with WRAP5:siRNA nanoparticles**

WRAP5:siRNA nanoparticles delivering siCYR61 and siCYR61’ (**A**,**B**) or siCTGF and siCTGF’ (**E**,**F**) induced a dose-dependent inhibition of CYR61 or CTGF in GIST-T1 cells after 48 h of incubation, as shown by Western blot quantification. No toxicity was observed respectively for siCYR61 and siCYR61’ (**C**,**D**) and siCTGF and siCTGF’ (**G**,**H**), as assessed by LDH assay. Controls include untreated (N.T.) and siNEG-treated cells. Data are presented as mean ± SD from N=4 independent experiments. Statistical analysis was performed using Kruskal-Wallis followed by Dunn’s multiple comparisons test *versus* siNEG. (**A**, **B**); ns >0.05, * <0.05, ** <0.01, *** <0.001.
